# Predicting base editing outcomes with an attention-based deep learning algorithm trained on high-throughput target library screens

**DOI:** 10.1101/2020.07.05.186544

**Authors:** Kim F. Marquart, Ahmed Allam, Sharan Janjuha, Anna Sintsova, Lukas Villiger, Nina Frey, Michael Krauthammer, Gerald Schwank

## Abstract

Base editors are chimeric ribonucleoprotein complexes consisting of a DNA-targeting CRISPR-Cas module and a single-stranded DNA deaminase. They enable conversion of C•G into T•A base pairs and vice versa on genomic DNA. While base editors have vast potential as genome editing tools for basic research and gene therapy, their application has been hampered by a broad variation in editing efficiencies on different genomic loci. Here we perform an extensive analysis of adenine- and cytosine base editors on thousands of lentivirally integrated genetic sequences and establish BE-DICT, an attention-based deep learning algorithm capable of predicting base editing outcomes with high accuracy. BE-DICT is a versatile tool that in principle can be trained on any novel base editor variant, facilitating the application of base editing for research and therapy.

## Introduction

Base editors are CRISPR/Cas-guided single-strand DNA deaminases. They enable precise genome editing by directly converting a targeted base into another, without the requirement of DNA double strand break formation and homology-directed repair from template DNA [1]. There are two major classes of base editors: cytosine base editors (CBEs) converting C•G into a T•A base pairs [2], and adenine base editors (ABEs) converting A•T into G•C base pairs [3]. The most commonly used base editors comprise a nickase (n) variant of SpCas9 to stimulate cellular mismatch repair, and have either the rat cytosine deaminase APOBEC1 or a laboratory-evolved *E. coli* adenine deaminase TadA fused to their N-termini. Both base editor classes convert target bases in a ∼5-nucleotide region within the protospacer target sequence, where the DNA strand that is not bound to the sgRNA becomes accessible to the deaminase [2], [3].

A major limitation of base editors is their broad variation in editing efficiencies across different target sequences. These can be influenced by several parameters, including the consensus sequence preference of the deaminase [4], and binding-efficiency of the sgRNA to the protospacer [5]. While low editing rates on a target locus that contains a single C or A may be circumvented by extending the exposure time to the base editor, undesired ‘bystander’ editing of additional C or A bases in the activity window generally requires optimization by experimental testing of alternative base editor constructs. Potentially successful strategies are i) exchanging the sgRNA to shift Cas9 binding up- or downstream, ii) using a base editor with a narrowed activity window [6], or iii) using a base editor with a deaminase that displays a different sequence preference (e.g. activation induced deaminase (AID) instead of rAPOBEC1 or ecTadA8e instead of ecTadA7.10) [7], [8]. Experimental screening with different available base editor and sgRNA combinations is however laborious and time-consuming, prompting us to establish a machine learning algorithm capable of predicting base editing outcomes of commonly used ABEs and CBEs on any given protospacer sequence *in silico*.

## Results and Discussion

### Generation of large datasets for ABEmax and CBE4max base editing via high-throughput screening with self-targeting libraries

To capture base editing outcomes of *Sp*Cas9 CBEs and ABEs across thousands of sites in a single experiment, we generated a pooled lentiviral library of constructs encoding unique 20-nt sgRNA spacers paired with their corresponding target sequences (20-nt protospacer and a downstream NGG PAM site) (Fig. 1a). Similar library designs have previously been used to analyze the influence of the sequence context on *Sp*Cas9 nuclease activity [9]–[12]. Our library included 18,946 target sites selected from randomly generated sequences, and 4,123 disease-associated human loci that can in theory be corrected with *Sp*Cas9 base editors (T-to-C or G-to-A mutations that harbor the PAM site 8-to-18 bp downstream of the target base) (Data file S1). An oligonucleotide library containing the sgRNAs and corresponding target sequences was synthesized in a pool, and cloned into a lentiviral backbone containing an upstream U6 promoter and a puromycin resistance cassette. HEK293T cells were then transduced at a 1000x coverage and a multiplicity of infection (MOI) of 0.5, ensuring that the majority of cells obtained a single integration. After puromycin selection, the cell pool was transfected with the two commonly used base editors ABE4max and CBE4max. ABEmax was evolved from the first reported adenine base editor ABE7.10 [13]. It contains the ecTadA7.10 deaminase domain N-terminally fused to n*Sp*Cas9, but has improved activity due to codon optimization and the use of bis-bpNLS domains. CBE4max is an enhanced version of the first reported cytidine base editor CBE3 with increased editing efficiency [13]. It is based on n*Sp*Cas9, N-terminally fused to rAPOBEC1 and C-terminally fused to two uracil glycosylase inhibitor (UGI) domains. After base editor transfection cells were cultured for four days, before genomic DNA was collected for high-throughput sequencing (HTS) to reveal editing outcomes at the target sites (Fig. 1b). In our analysis we first filtered for a set of target sites that were sampled with at least 1,000 reads across the three experimental replicates and obtained at least 1.5% reads with base editor specific nucleotide conversions. This led to the identification of a subset of 9,011 sites for CBE4max and 6,641 sites for ABEmax (Data file S1). In line with previous studies, we found that base editing efficiencies (defined here as the fraction of mutant reads over all sampled reads of a target site) are lower when the sgRNA is expressed from a lentivirally integrated cassette compared to a transfected plasmid (median overall editing efficiency ABEmax= 2.2%, CBE4max= 2.4%)(Fig. S1) [14], [15]. The editing outcomes and frequencies were nevertheless consistent between the three experimental replicates (median Pearson’s *R* = 0.89 (ABEmax) and 0.95 (CBE4max)) (Fig. S2), indicating robust base editor activity and robust sampling of WT and edited reads. In addition, the obtained activity windows of ABEmax and CBE4max were similar to previous studies, with maximum editing at position 6 (Fig. 1c, d). Analysis of the trinucleotide sequence context moreover confirmed previous studies, which showed that ecTadA7.10 and rAPOBEC1 have a preference for editing at bases that are preceded by T (Fig. 1e, f) [16], [17]. Interestingly, for ecTadA7.10 we also observed an aversion for an upstream A and preference for a downstream C, while in case of rAPOBEC1 we observed an aversion for upstream C or G and downstream C (Fig. 1e, f).

**Figure 1:**
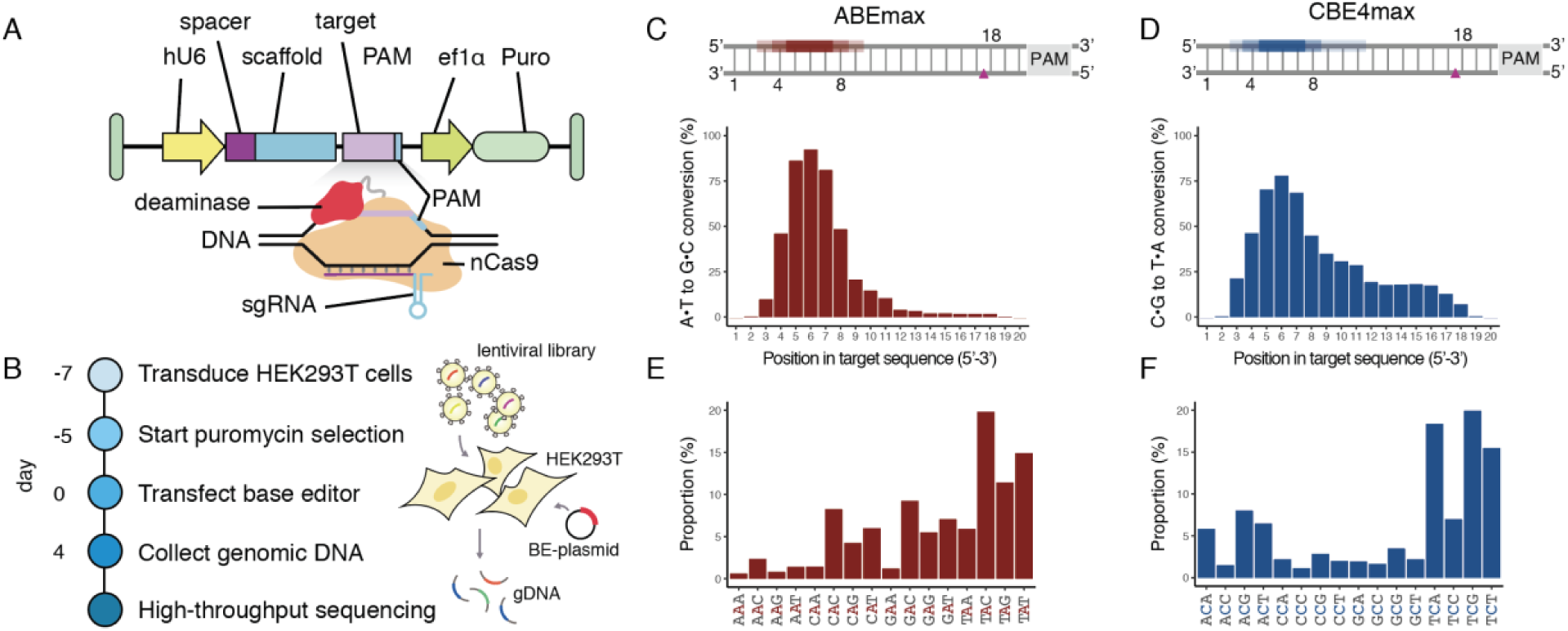
A high-throughput platform for assessing base editor activities. (**A**) Design of the self-targeting library: The lentiviral library contains the sgRNA expression cassette and the target locus on the same DNA molecule. The sgRNA (spacer and scaffold) is transcribed under the control of a U6 promoter and is designed to direct the base editor (nCas9-deaminase fusion) to the 20-nt sequence upstream of the protospacer adjacent motif (PAM). hU6, human U6 promoter; ef1α, elongation factor 1α promoter; nCas9, nickase Cas9; sgRNA, single-guide RNA; Puro, puromycin selection marker. (**B**) Overview of library screening. (**C**,**D**) Distribution of edited bases for (**C**) ABEmax and (**D**) CBE4max. The plot shows the proportion of A-to-G or C-to-T base conversions ≥1.5% at each position across the protospacer target sequence. The schematic (top) illustrates the favored activity window of the respective deaminase. The nCas9 induced nick is indicated as a triangle. (**E**,**F**) Proportion of the different tri-nucleotide motifs at bases edited by ABEmax and CBE4max ≥1.5%.

### Development of BE-DICT, an attention-based deep learning model predicting base editing outcomes

Potentially predictive features that influence CRISPR/Cas9 sgRNA activity, such as the GC content and minimum Gibbs free energy of the sgRNA, did not influence base editing rates (Suppl. Fig. 3a, b). Moreover, DeepSpCas9 [10], a deep learning tool designed to predict cutting efficiencies of Cas9, could not be repurposed to forecast base editing rates (Pearson’s R for editing of an A or C anywhere in the protospacer is -0.07 for ABEmax and 0.04 for CBE4max) (Fig. S4). This prompted us to utilize the comprehensive base editing data generated in the ABEmax and CBE4max target library screens for designing and training a machine learning model capable of predicting base editing outcomes at any given target site. We established BE-DICT (Base Editing preDICTion via attention-based deep learning), an attention-based deep learning algorithm that models and interprets dependencies of base editing on the protospacer target sequence. The model is based on multi-head self-attention inspired by the Transformer encoder architecture [18]. It takes a sequence of nucleotides of the protospacer as input and computes the probability of editing for each target nucleotide as output (Fig. 2a). Formal description of the model and the different computations involved are reported in the Supplementary Materials. In short, BE-DICT assigns a weight (attention-score) to each base within the protospacer (i.e. learned fixed-vector representation), which is used to calculate the probability with which a target base will be converted from A-to-G or C-to-T (probability score). As a binary input mode was chosen where all bases ≥ 1.5% editing (C-to-T or A-to-G) were classified as edited and bases < 1.5% editing were classified as unedited, the probability score reflects the likelihood with which a target base is classified as edited or not. To train and test the model, we used data from 11,246 and 8,381 target sites of the CBE4max and ABEmax screen, respectively (Data file S1). For model training we used ∼80% of the dataset, and performed stratified random splits for the rest of the sequences to generate an equal ratio (1:1) between test and validation datasets. We repeated this process five times (denoted by runs), in which we trained and evaluated a model for every base editor separately for each run. BE-DICT performance was evaluated using area under the receiver operating characteristic curve (AUC), and area under the precision recall curve (AUPR). An average AUC equal to 0.960 ± 0.0027 and an average AUPR equal to 0.849 ± 0.0039 (mean ± s.d.) was achieved for ABEmax, and an average AUC equal to 0.9178 ± 0.0044 and an average AUPR of 0.719 ± 0.0147 was achieved for CBE4max (Fig 2b, c). Notably, at positions within the activity window where we have a balanced distribution of edited vs. unedited substitute bases, BE-DICT performed with significantly higher accuracy than a per position majority class predictor (a baseline that models the observed target nucleotides and their conversions at each position across all target sites in the training data as a Bernoulli trial and uses maximum likelihood estimation for computing the probability of editing success) (Fig. 2c, Fig. S5).

**Figure 2:**
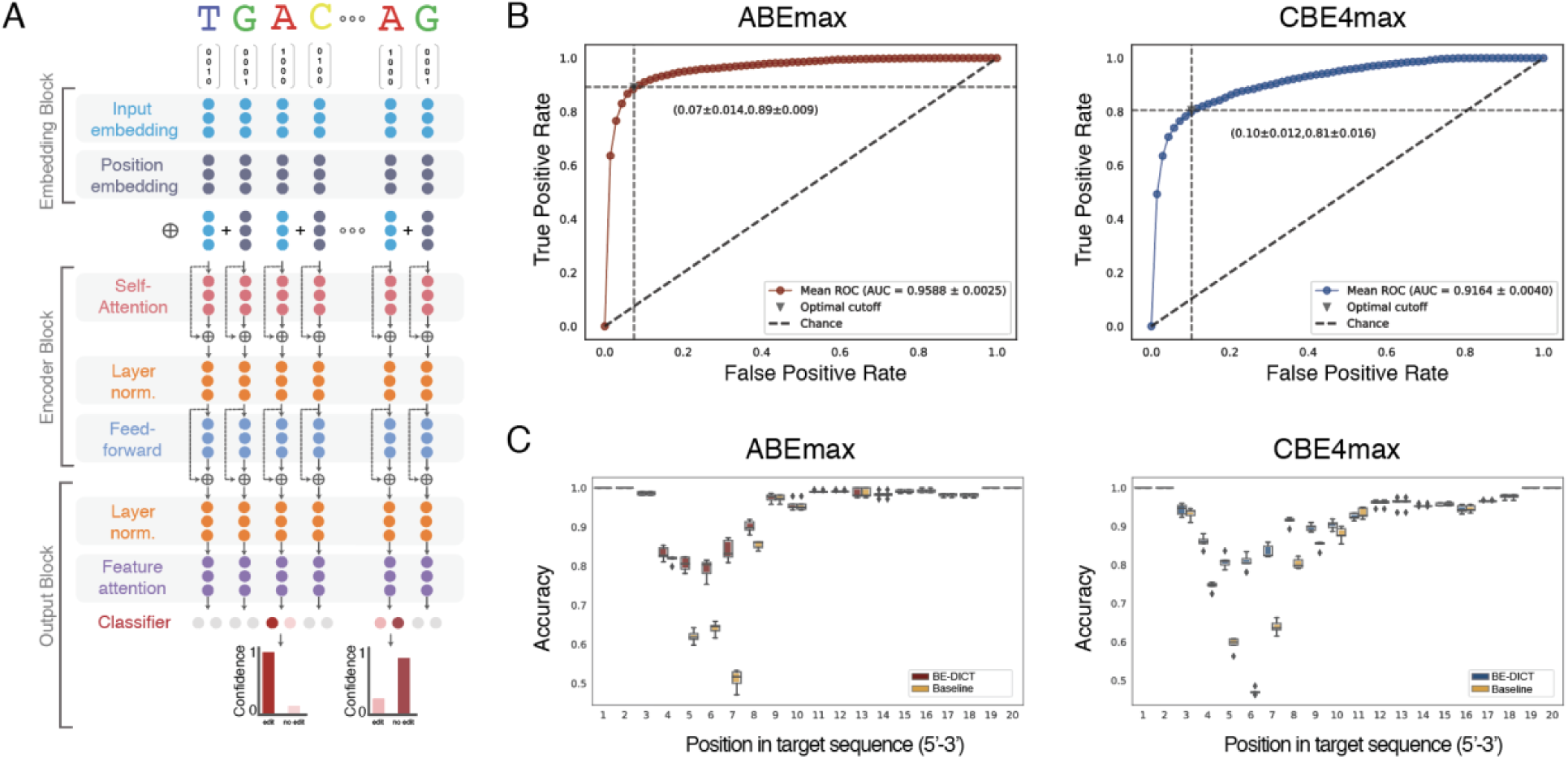
BE-DICT: A machine learning model for predicting base editing outcomes. (**A**) Design of an attention-based deep learning algorithm. Given a target sequence, the model can be queried with a confidence score to predict the chance of target base conversion. The model has three main blocks: (1) An embedding block that embeds both the nucleotide and its corresponding position from one-hot encoded representation to a dense fixed-length vector representation. (2) An encoder block that contains (a) a self-attention layer (with multi-head support), (b) layer normalization [26] and residual connections [27] and (c) a feed-forward network. (3) An output block that contains (a) a position attention layer, and (b) a classifier layer. (**B**) The average AUC achieved across 5 runs (interpolated) for models trained on data from ABEmax (left) and CBE4max (right). (**C**) Box plot of per-position accuracy of the trained models across 5 runs for both BE-DICT ABEmax and CBE4max editors in comparison to the accuracy of majority class baseline predictor.

**Figure 3:**
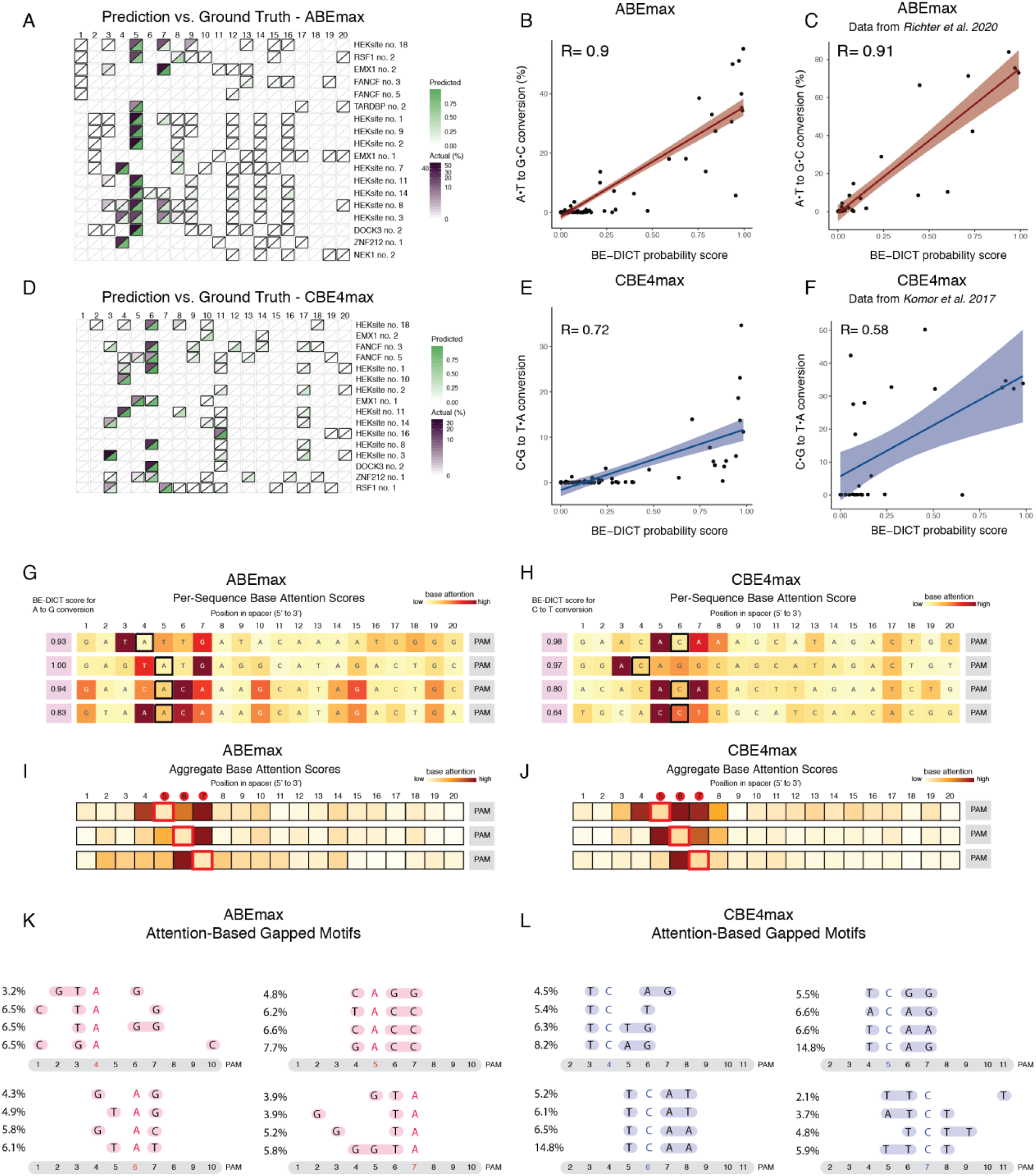
BE-DICT accurately predicts base editing activities on genomic loci and predominantly puts attention on nucleotides flanking the target base. Protospacers with at least two substrate nucleotides were targeted by either ABEmax or CBE4max. (**A**) Heatmap shows the BE-DICT prediction values (green) and the A-to-G editing percentage (purple) for sequences targeted with ABEmax. (**B**) Correlation between BE-DICT probability score and the A-to-G editing percentage for sequences shown in A. (**C**) Correlation between BE-DICT probability score and the A-to-G editing percentage for sequences from *Richter et al*. [8] (**D**) Heatmap shows the BE-DICT probability scores (green) and the editing percentage (purple) for sequences targeted with CBE4max. (**E**) Correlation between the BE-DICT probability score and the C-to-T editing percentage for sequences shown in E. (**F**) Correlation between BE-DICT probability score and the C-to-T editing percentage for sequences from *Komor et al*. [19] (**G**,**H**) BE-DICT attention scores for the base indicated in bold and the respective prediction value for editing for sequences targeted with ABEmax and CBE4max. Attention weight is color indicated from light to dark. (**I**,**J**) Mean-aggregate attention scores of target sequences predicted to be edited by BE-DICT at positions 4, 5 and 6 for ABEmax (n = 1,226) and CBE4max (n = 913). (**K**,**L**) Most frequently occurring gapped-motifs based on mean-aggregate attention scores of the target sequence. The target base is highlighted in red (ABEmax) and blue (CBE4max). The percentages (left) indicate the frequency of occurrence of the respective gapped-motifs.

### BE-DICT can be utilized to predict editing efficiencies at endogenous loci and predominantly puts attention to bases flanking the target base

Base editing at endogenous loci may also be affected by protospacer sequence independent factors, such as chromatin accessibility. We therefore next tested the accuracy of BE-DICT in predicting base editing outcomes at 18 separate endogenous genomic loci for ABEmax, and 16 endogenous genomic loci for CBE4max. HEK293T cells were co-transfected with plasmids expressing the sgRNA and base editor, and genomic DNA was isolated after four days for targeted amplicon HTS analysis. Across all tested loci we observed a strong correlation between experimental editing rates and the BE-DICT probability score (Pearson’s *R* = 0.9 for ABEmax and 0.72 for CBE4max; Fig. 3a, b, d, e; Data file S2). Further validating our model, BE-DICT also accurately predicted base editing efficiencies from previously published experiments (Pearson *R* = 0.91 for ABEmax and 0.58 for CBE4max; Fig. 3c, f; Fig. S6; Data file S2) [8], [19]. These results demonstrate that the BE-DICT probability score can be used as a proxy to predict ABEmax and CBE4max editing efficiencies with high accuracy. One benefit of the attention-based BE-DICT model is that the influence of each position within the protospacer on the decision whether a target base is edited or not can be inspected. These attention scores provide a proxy for identifying relevant motifs and sequence contexts for editing outcomes. As expected, for both base editors BE-DICT attention was most often focused on the bases flanking the target base (Fig. 3g-h). In addition, we found that base attention patterns were dependent on the position of the target base within the protospacer sequence (Fig. 3i-j), and sometimes consisted of complex gapped motifs rather than consecutive bases (Fig. 3 k-l). These observations underscore the necessity of using machine learning algorithms for predicting base editing outcomes from protospacer sequences.

### BE-DICT can be trained for predicting base editing outcomes of ABE8e and Target-AID

For loci where the sequence preference of rAPOBEC1 or ecTadA7.10 prevents efficient editing, the use of base editors that employ different deaminases might be successful. We therefore next performed the target library screen with Target-AID, an alternative CBE that is based on the activation-induced cytidine deaminase (AID) ortholog *Pm*CDA1 [7], and ABE8e, an alternative ABE that is based on a further evolved ecTadA deaminase with increased activity [8]. HEK293T cells containing the integrated target sites were transfected with plasmids expressing the base editor and incubated for four days before extraction of genomic DNA and HTS analysis. Editing rates were consistent between the three experimental replicates (median Pearson’s *R* = 0.84 for ABE8e and 0.95 for Target-AID; Fig. S7), and filtering target sites for ≥ 1000 reads and editing rates ≥ 1.5 % resulted in 4,088 sites for Target-AID and 4,184 sites for ABE8e (Fig. S8; Data file S1). Confirming the observation by Richter et al. [8], we found that ABE8e has a broader activity window than ABE7.10max (Fig. 4a). Trinucleotide consensus motif analysis moreover revealed that ABE8e displayed a reduced sequence preference, although editing of bases that were preceded by an A were still largely disfavored (Fig. 4b). In addition, for Target-AID the activity window was extended compared to CBE4max (Fig. 4c), and while *Pm*CDA1 lacked the requirement of a preceding T for efficient editing, motifs where the targeted base is followed by a C were slightly disfavored (Fig. 4d). When we next trained BE-DICT with ABE8e and Target-AID screening datasets, we observed strong performance in predicting editing outcomes. We obtained an AUC = 0.9317 ± 0.0063 for ABE8e and 0.858 ± 0.0063 for Target-AID, and an AUPR= 0.775 ± 0.0220 for ABE8e and 0.570 ± 0.027 for Target-AID (Fig. 4e, f). Similar to ABEmax and CBE4max, both models achieved significantly higher accuracy compared to the per position majority class predictor at positions within the activity window, where we had a balanced distribution of edited vs. unedited substrate bases (Fig. 4g, h; Fig. S9). Importantly, the generated editing probability also correlated well with the editing efficiencies of ABE8e and Target-AID at multiple endogenous loci (Pearson’s *R* = 0.6 - 0.86; Fig. 4i-n). Together these results suggest that BE-DICT is a versatile tool, which can be trained to predict editing efficiencies of various different base editor variants.

**Figure 4:**
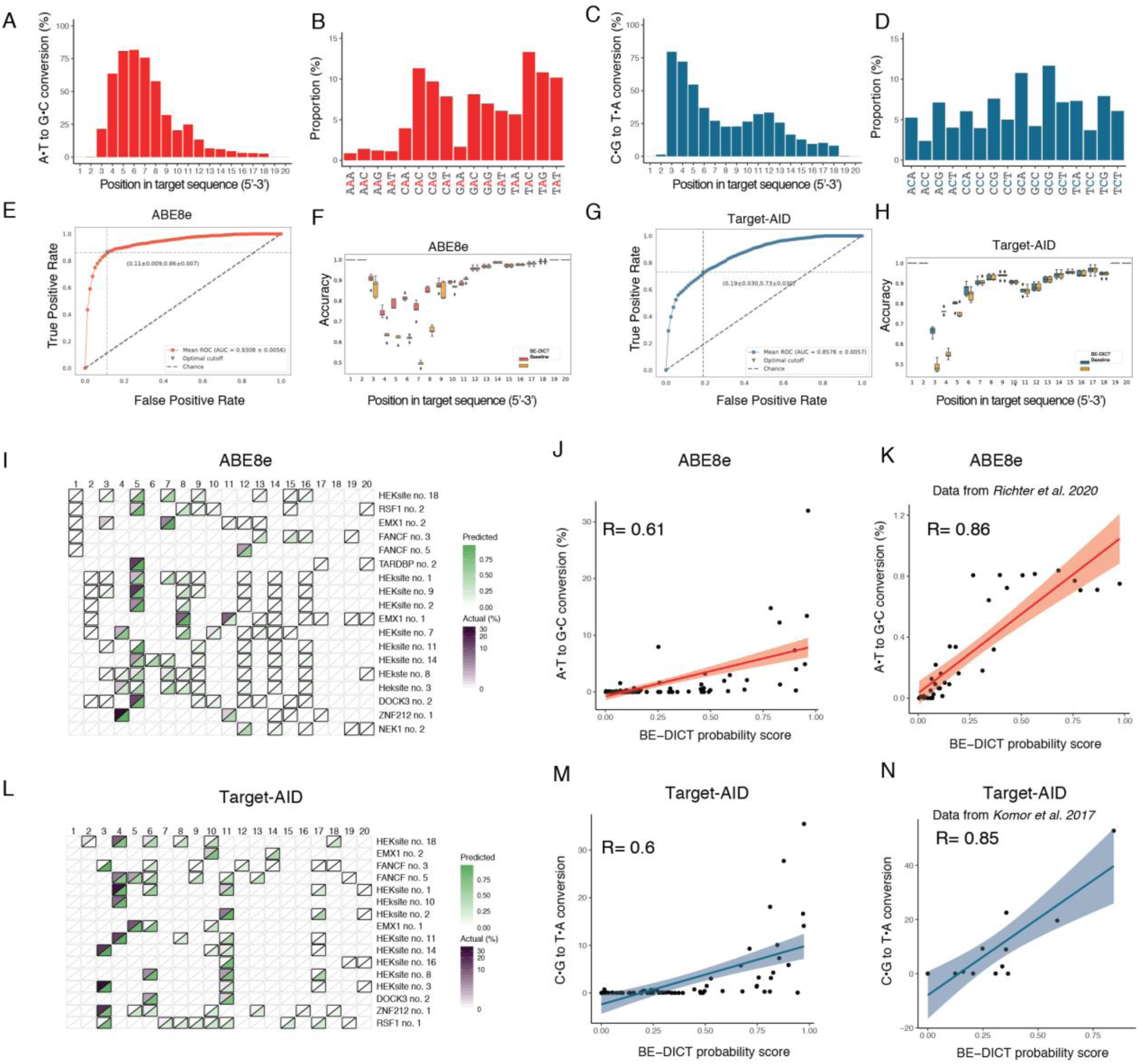
Expanding the capability of BE-DICT to ABE8e and Target-AID base editors. (**A**) Relative distribution of A-to-G edited bases ≥1.5% and (**B**) proportion of tri-nucleotide sequence motifs of A-to-G edited bases ≥1.5% for ABE8e. (**C**) Relative distribution of C-to-T edited bases ≥1.5% and (**D**) proportion of tri-nucleotide sequence motifs of C-to-T edited bases ≥1.5% for Target-AID. Average AUC across 5 runs (interpolated) of the BE-DICT model trained on ABE8e (**E**) and Target-AID (**G**), and per-position accuracy of both base editors (**F**,**H**) in comparison to the accuracy of a majority class baseline predictor. (**I**) Heatmap shows BE-DICT probability scores (green) and the editing percentage (purple) for sequences targeted with ABE8e. (**J**) Correlation between the BE-DICT probability score and the A-to-G editing percentage for sequences shown in (I). (**K**) Correlation between the BE-DICT probability score and the A-to-G editing percentage for sequences from *Richter et al*. [8] (**L**) Heatmap shows the BE-DICT prediction values (green) and the editing percentage (purple) for sequences targeted with Target-AID. (**M**) Correlation between the BE-DICT probability score and the C-to-T editing percentage for sequences shown in (L). (**N**) Correlation between BE-DICT probability score and the C-to-T editing percentage for sequences from *Komor et al*. [19].

### Comparison between BE-DICT and BE-Hive

At the time we prepared this manuscript, a deep conditional autoregressive machine learning model (BE-Hive) capable of predicting editing outcomes and efficiencies of several base editors including ABEmax and CBE4max was published [20]. To compare the performance of both models, we applied BE-Hive to the same set of endogenous loci tested in our study (Fig. 3b-f, 4h-l). Overall BE-Hive achieved Pearson’s *R* values for predicting base editing efficiencies in the range of BE-DICT (0.77-0.82 for ABEmax and 0.46-0.85 for CBE4max; Fig. S11). Interestingly, both models were better in predicting ABEmax editing efficiencies compared to CBE4max editing efficiencies, which is in line with our observation that BE-Hive and BE-DICT prediction values show a stronger correlation for ABEmax (Pearson’s *R* = 0.88-0.89) than for CBE4max (Pearson’s *R* = 0.55-0.67) (Fig. S11). Notably, the BE-hive model considers 4^N^ (the space of all possible base conversion combinations), where N is the number of nucleotides in the target site. As a consequence, a heuristic approach was used to limit the search space in this exponential setting, providing probabilities for most but not all possible editing combinations. In contrast, BE-DICT models the marginal probability at each position and thus the complexity increases only linearly with the nucleotide number by further scaling the self-attention layer to O(n) complexity [21], enabling the model to consider a wider sequence context beyond the protospacer target site.

In summary, in this study we performed high-throughput base editing experiments with four commonly used base editors, and employed these data sets as input to establish BE-DICT, a deep learning model capable of accurately predicting editing outcomes. As the model is attention based, it can readily be used to extract motifs that disfavor editing with currently available deaminases, potentially guiding researchers towards rational development of novel base editor variants. BE-DICT was trained on datasets for ABEmax, CBE4max, ABE8e and Target-AID, and will be freely accessible at https://be-dict.org. The algorithm is versatile, and in the future could also be adopted to various other base editor variants, allowing researchers a priori selection of an optimal base editor for their target locus.

## Materials and Methods

### Oligonucleotide-library design

The custom oligonucleotide pool containing pairs of sgRNA and corresponding target sequences was purchased from Twist Bioscience. Designed oligonucleotides include the following elements: The G/20N spacer and SpCas9 gRNA scaffold, a 6-nucleotide randomized barcode, the corresponding target locus containing the PAM and a second 6-nucleotide randomized barcode (Supplementary material). The library includes 18,946 random sequences and 4,123 disease loci theoretically targetable using base editors. The disease loci were selected from the NCBI ClinVar [22] database (filtered as pathogenic, clinical and monogenic; accessed on May, 2019) using the following criteria: The sequences should contain a target A or C in the editing window and contain an NGG PAM 8-to-18 bases away from the target base.

### Plasmid-library preparation

The plasmid library containing the sgRNA and the corresponding target sequence was prepared using a one-step cloning process to prevent uncoupling of the sgRNA- and target sequence. The oligonucleotide pool was PCR-amplified in 10 cycles (Primers stated in Supplemental Information) and KAPA® HiFi HotStart Polymerase (Roche) following the manufacturer’s instructions. The resulting amplicons were then purified using 0.8x volumes of paramagnetic AMPure XP beads (Beckman Coulter) following the manufacturer’s instructions for PCR cleanup. We digested the Lenti-gRNA-Puro plasmid with BsmBI restriction enzyme (New England Biolabs, NEB) for 12 h at 55 °C. Lenti-gRNA-Puro was a gift from Hyongbum Kim (Addgene no. 84752). After digestion, the plasmid was treated with calf intestinal alkaline phosphatase (NEB) for 30 min at 37 °C and gel purified with a NucleoSpin Gel and PCR Clean-up Mini kit (Macherey-Nagel). The oligo-pool amplicons were assembled into the linearized Lenti-gRNA-Puro plasmid using NEBuilder HiFi DNA Assembly Master Mix (NEB) for 1 h at 50 °C. The product was precipitated by adding one volume of Isopropanol (99%), 0.01 volumes of GlycoBlue coprecipitant (Invitrogen) and 0.02 volumes of 5M NaCl solution. The mix was vortexed for 10 seconds and incubated at room temperature for 15 min. followed by 15 min. of centrifugation (15,000 x g). The supernatant was discarded and replaced by two volumes of ice-cold ethanol (80%). Ethanol was removed immediately and the pellet was air-dried for 1 min. The pellet was dissolved in TE buffer (10 mM Tris, 0.1 mM EDTA) and incubated at 55 °C for 10 min. In 12 transformation replicates, 2 µL of plasmid library were transformed per 50 µL of electrocompetent cells (ElectroMAX Stabl4, Invitrogen) using a Gene Pulser II device (Bio-Rad). Transformed cells were recovered in S.O.C media (from ElectroMAX Stabl4 kit) for 1h and spread on Luria–Bertani agar plates (245×245 mm, Thermo Fisher Scientific) containing 100 µg/mL ampicillin. After incubation at 30 °C for 14 h, the colonies were scraped and plasmids were purified using a Plasmid Maxiprep kit (Qiagen).

### Cell culture

HEK293T (ATCC CRL-3216) were maintained in DMEM plus GlutaMax (Thermo Fisher Scientific), supplemented with 10% (vol/vol) Fetal Bovine Serum (FBS, Sigma-Aldrich) and 1× penicillin–streptomycin (Thermo Fisher Scientific) at 37 °C and 5% CO2. Cells were maintained at confluency below 90% and passaged every 2-3 days. Cells were discarded after 15 consecutive passages.

### Packaging of guide RNA library into lentivirus

Transfection in T225 cell culture flasks was conducted as follows: 3.4 µg pCMV-VSV-G (lentiviral helper plasmid; addgene plasmid no. 8454, a gift from B. Weinberg), 6.8 µg psPAX2 (lentiviral helper plasmid; addgene plasmid no. 12260, a gift from D. Trono) and 13.6 µg target library plasmid were mixed in 651 µL Opti-MEM (serum free, Opti-MEM (Thermo Fisher Scientific)), supplied with 195 µL of polyethyleneimine (PEI, 1 mg/mL), vortexed for 10 sec and incubated for 10 min. 25 mL of DMEM was added to the transfection mix and gently pipetted to the cells at approximately 70% confluency. Medium was changed 1 day after transfection. 2 days later, supernatant-containing lentiviral particles were harvested and filtered using a Filtropur S 0.4 (Sarstedt) filter. The virus suspension was ultracentrifuged (20,000 x g) for 2 h. Aliquots were frozen at −80 °C until use.

### Pooled base editor screens

T175 cell culture flasks were seeded with HEK293T cells and cultured to reach 70-80% confluence. 10 µg/mL polybrene was added to the media and the gRNA-pool lentivirus was transduced at a multiplicity of infection (MOI) of 0.5 and a calculated coverage of 1000 cells per gRNA. One day after transduction, cells were supplied with fresh media with 2.5 µg/mL Puromycin. After 9 days of puromycin selection, the respective base editor plasmids were transfected using 50 µg of plasmid DNA in a 1:3 DNA:PEI ratio per T175 flask. 4 days later, cells were detached and genomic DNA was extracted using a Blood & Cell Culture DNA Maxi kit (Qiagen) according to the manufacturer’s instructions. Base editor plasmids pCMV-ABEmax-P2A-GFP (Plasmid no. 112101), pCMV-BE4max-P2A-GFP (Plasmid no. 112099) and pCMV-ABE8e (Plasmid no. 138489) were a gift from David Liu. Target-AID (pRZ762, Plasmid no. 131300) was a gift from Keith Joung.

### Guide RNA cloning

The vector backbone (lentiGuide-Puro, Plasmid no. 52963) was digested with Esp3I (NEB) and treated with rSAP (NEB) at 37 °C for 3 hours and gel-purified on a 0.5 % agarose gel. For (Suppl. Information 1) sgRNA phospho-annealing, 1 µL of sgRNA top and bottom strand oligonucleotide (100 µM each), 1 µL 10x T4 DNA Ligase Buffer, 1 µL T4 PNK (NEB) and 6 µL H2O were mixed and incubated in a thermocycler (BioRad) using the following program: 37 °C for 30 min, 95 °C for 5 min, ramp down to 25 °C at a rate of 5 °C/min. The annealed oligonucleotides were diluted 1:100 in H2O and ligated into the vector backbone using 50 ng digested lentiGuide-Puro plasmid, 1 µL annealed oligonucleotide, 1 µL 10x Ligase Buffer (NEB) and 1 µL T4 DNA Ligase (NEB) in a 10 µL reaction (filled up to total volume with H2O). The ligation mix was incubated at room temperature for 3 h and transformed into NEB Stable Competent E. coli (C3040H) following the manufacturer’s instructions. Correct assembly of the sgRNA into the backbone was confirmed by SANGER-Sequencing (Microsynth) and plasmids were isolated using a GeneJET Plasmid Miniprep Kit (Thermo Fisher Scientific) following the manufacturer’s instructions.

### Arrayed sgRNA transfections

For base editor DNA on-target experiments HEK293T cells were seeded into 96-well flat-bottom cell culture plates (Corning), transfected 24 h after seeding with 150 ng of base editor and 50 ng of gRNA expression plasmid and 0.5 µl of Lipofectamine 2000 (Invitrogen) per well. One day later, medium was removed and cells were detached using one drop of TrypLE (Gibco) per well, resuspended in fresh medium containing 2.5 µg/uL puromycin and plated again into 96-well flat-bottom cell culture plates. Cells were detached 4 days after transfection and pelleted by centrifugation. To obtain genomic DNA, cells were resuspended in 30 µL 1x PBS and 10 µL of lysis buffer (4x Lysis Buffer: 10 mM Tris-HCL at pH8, 2% Triton X, 1 mM EDTA and 1% freshly added Proteinase K (Qiagen)) was added to the cell suspension. The lysis was performed in a thermocycler (Bio-Rad) using the following program: 60 °C, 60 min; 95 °C, 10 min; 4 °C, hold. The lysate was diluted to a final volume of 100 µL using nuclease-free water and 1 µL of each lysate was used for the subsequent PCR.

### Library preparation for targeted amplicon sequencing of DNA

Next-generation sequencing (NGS) preparation of DNA was performed as previously described [23]. In short, the first PCR was performed to amplify genomic sites of interest with primers containing Illumina forward and reverse adaptor sequences (see Supplemental material for primers and amplicons used in this study). To cope with high DNA input used for pooled screens, the Herculase II Fusion DNA Polymerase (Agilent) was used according to the manufacturer’s instructions. For all other NGS-PCRs on genomic DNA and the second NGS-PCR, the NEBNext High-Fidelity 2× PCR Master Mix (NEB) was used according to the manufacturer’s instructions. In brief, 0.96 mg of genomic DNA per replicate of the pooled gRNA screen was amplified in 24 cycles for the first PCR using 10 µg gDNA input in 100 µL reactions. For arrayed gRNA experiments, 1 µL of the cell lysate per replicate was used in a 12.5 µL PCR reaction. The first PCR products were cleaned with paramagnetic beads, then the second PCR was performed to add barcodes with primers containing unique sets of p5/p7 Illumina barcodes (analogous to TruSeq indexes). The second PCR products were again cleaned with paramagnetic beads. The final pool was quantified on the Qubit 4 (Invitrogen) instrument. Pooled sgRNA screens were sequenced single-end on the Illumina NovaSeq 6000 machine using a S1 Reagent Kit (100 cycles). Arrayed gRNA experiments were sequenced paired-end (2 × 150) on the Illumina MiSeq machine using a MiSeq Reagent Kit v2 Nano.

### High-throughput sequencing analysis

Fastq reads obtained from deep sequencing were trimmed upto the guide sequence by removing the Illumina adapters and the plasmid scaffold sequences using Cutadapt v2.2 [24]. The trimmed reads were then mapped using bowtie2 v2.3.5.1 [25] with default parameters to a reference consisting of target sequences making up the library. Mismatches identified from the aligned reads were filtered for C-to-T (in case of CBE) or A-to-G (in case of ABE) conversions in the protospacer. The sequences were further filtered for a read depth of at least 1,000. Only the sequences passing this filtering criteria were used for further analysis.

### Statistical Analysis

Editing percentages and correlation analysis using Pearson’s R was calculated using R 3.5.2 and Python3.

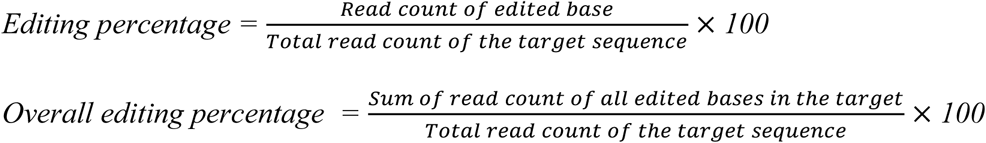

DeltaG was calculated using the online resource at http://unafold.rna.albany.edu/.

## H2: Supplementary Materials

Fig. S1: Editing rates in the self-targeting pooled library for ABEmax and CBE4max.

Fig. S2: Correlations between the results of technical replicates of the high-throughput screening experiments.

Fig. S3: GC-content (%) and Gibbs free energy (ΔG) for ABEmax and CBE4max in edited sequences and non-edited sequences.

Fig. S4: DeepSpCas9 score prediction versus base editing efficiency.

Fig. S5: Per position target base conversion (A-to-G; C-to-T) in the Training and Validation/Test set of the pooled ABEmax and CBE4max screen used for BE-DICT.

Fig. S6: Comparison of predicted (BE-DICT) versus published base editing activities on genomic loci in HEK293T cells for ABEmax (compare Richter et al. 2020 [8]) and CBE4max (compare Komor et al. 2017 [19]).

Fig. S7: Correlations between the results of technical replicates of the high-throughput screening experiments.

Fig. S8: Editing rates in the self-targeting pooled library for ABE8e and Target-AID screens.

Fig. S9: Per position substrate base conversion (A-to-G; C-to-T) in the Training and Validation/Test set of the pooled ABE8e and Target-AID screen used for BE-DICT.

Fig. S10: Comparison of predicted (BE-DICT) versus published base editing activities on genomic loci for ABE8e (compare Richter et al. 2020 [8]) and Target-AID (compare Komor et al. 2017 [19]).

Fig. S11: Comparison of BE-Hive (Arbab et al. 2020 [20]) and BE-DICT performance.

Data file S1: BE-DICT train and test dataset.

Data file S2: HTS analysis of arrayed sgRNA transfections.

## General

We thank the Functional Genomics Center Zurich (FGCZ) for their help and support.

## Funding

This work was supported by the SNF (310030_185293). S.J. holds an EMBO Long-Term Fellowship (ALTF 873-2019), and K.F.M holds a PHRT iDoc Fellowship (PHRT_324).

## Author Contributions

K.F.M. designed the study, performed experiments, and analyzed data. A.A. and A.S. designed and generated the machine learning algorithm BE-DICT and analyzed data. S.J. designed the study and analyzed the high-throughput sequencing data. L.V. and N.F. performed experiments. M.K. and G.S. designed and supervised the research and wrote the manuscript. K.F.M., A.A., S.J., and A.S. edited the manuscript. All authors approved the final version of the manuscript.

## Competing interests

The authors declare no competing interests.

## Data and materials availability

We have provided the data sets used in this study as Data files 1-2. DNA-sequencing data are deposited under accession number GEO: (tba).

We have made the source code for BE-DICT and the custom Python scripts used to train and evaluate the models available on Github at https://github.com/uzh-dqbm-cmi/crispr. The web application for BE-DICT, which predicts the base editing patterns in the DNA sequence will be made available at https://be-dict.org.

## Supplementary Materials

**Supplemental figure 1:**
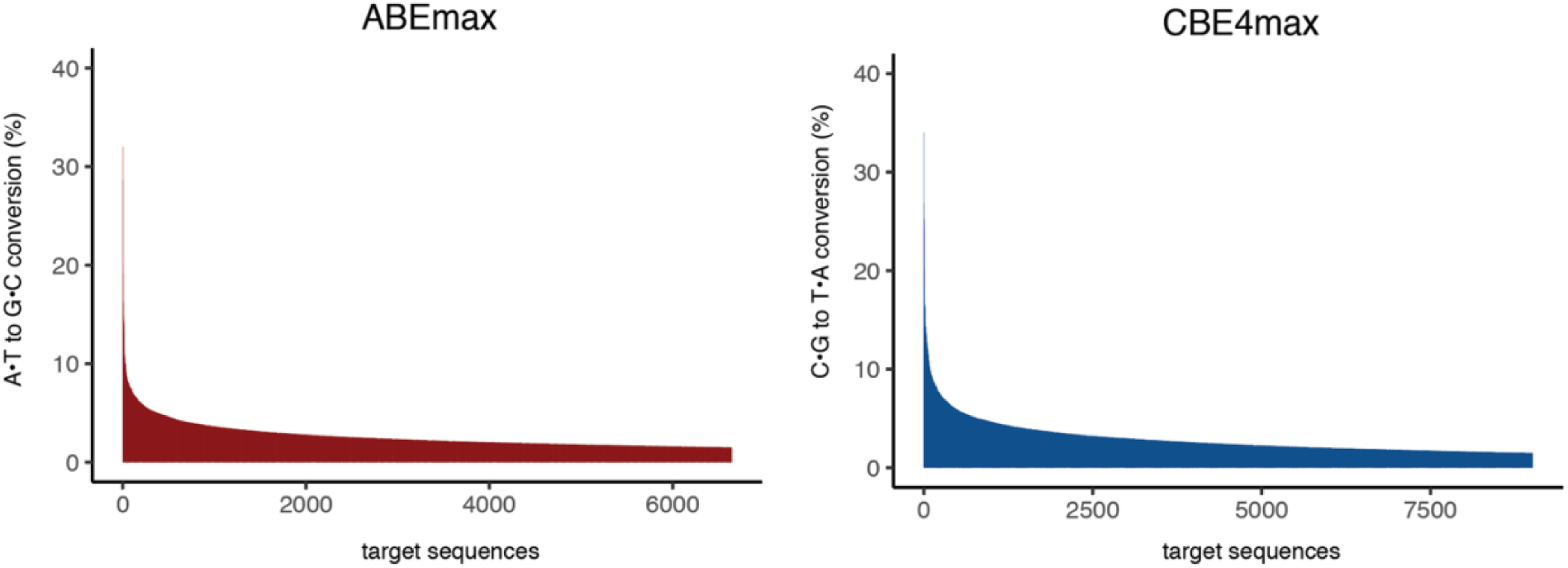
Editing rates in the self-targeting pooled library for ABEmax and CBE4max. Percentage of overall editing efficiency scores per target sequence for ABEmax and CBE4max.

**Supplemental figure 2:**
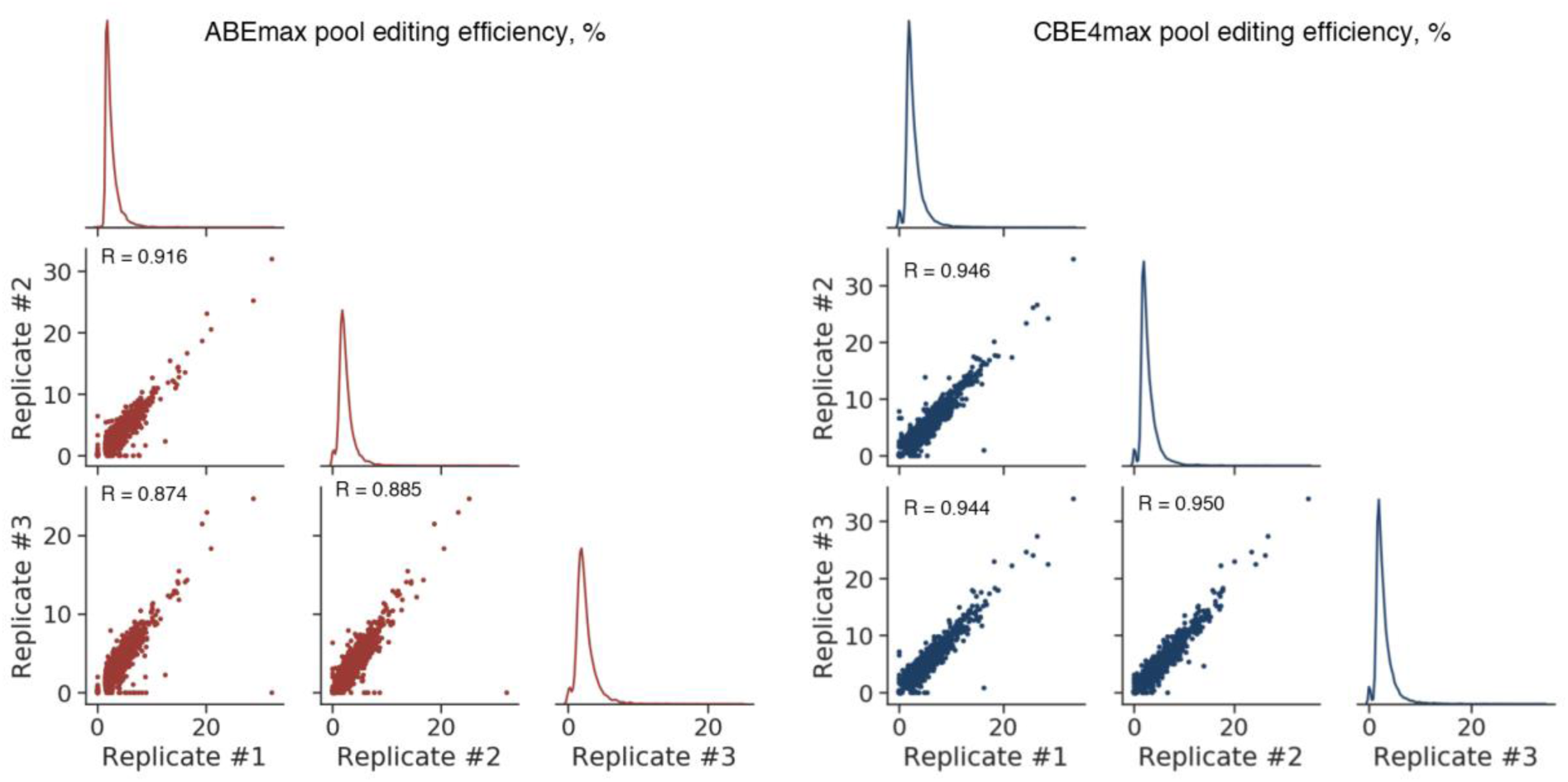
Correlations between the results of technical replicates of the high-throughput screening experiments. Scatter plots show the correlation between editing efficiencies for all sequences that were edited in at least one of the three technical replicates (with editing efficiency ≥1.5%) from experiments of library-cells treated with ABEmax and CBE4max.

**Supplemental figure 3:**
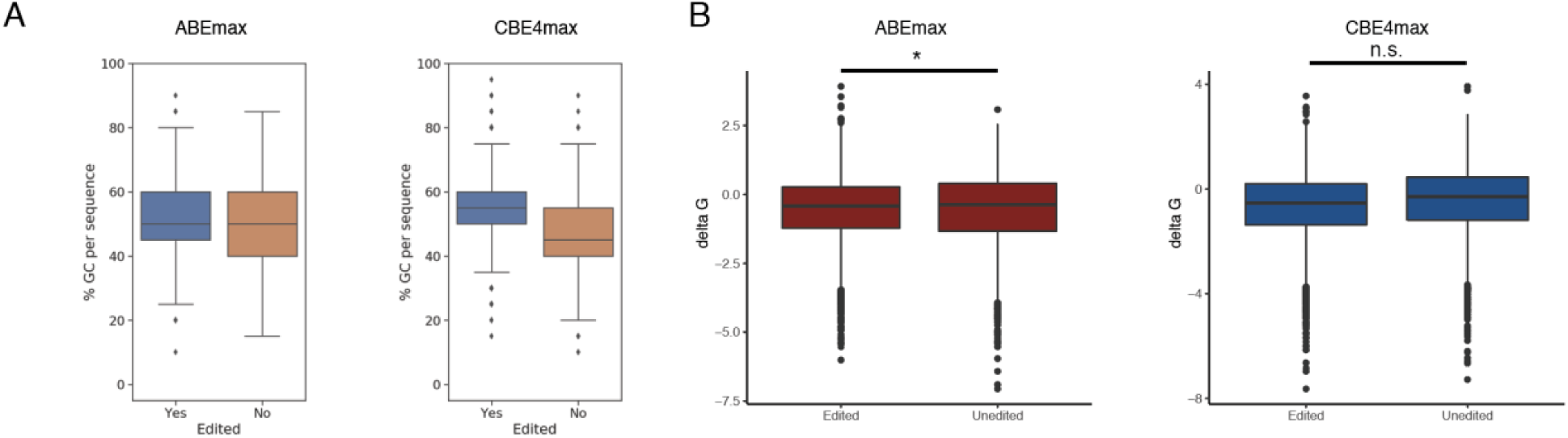
GC-content (%) and Gibbs free energy (ΔG) for ABEmax and CBE4max in edited sequences and non-edited sequences. Boxplots show the distribution of GC content (A) and delta G values (B) of all edited and unedited target sequences for ABEmax and CBE4max (*, p < 0.05; n.s. not-significant).

**Supplemental figure 4:**
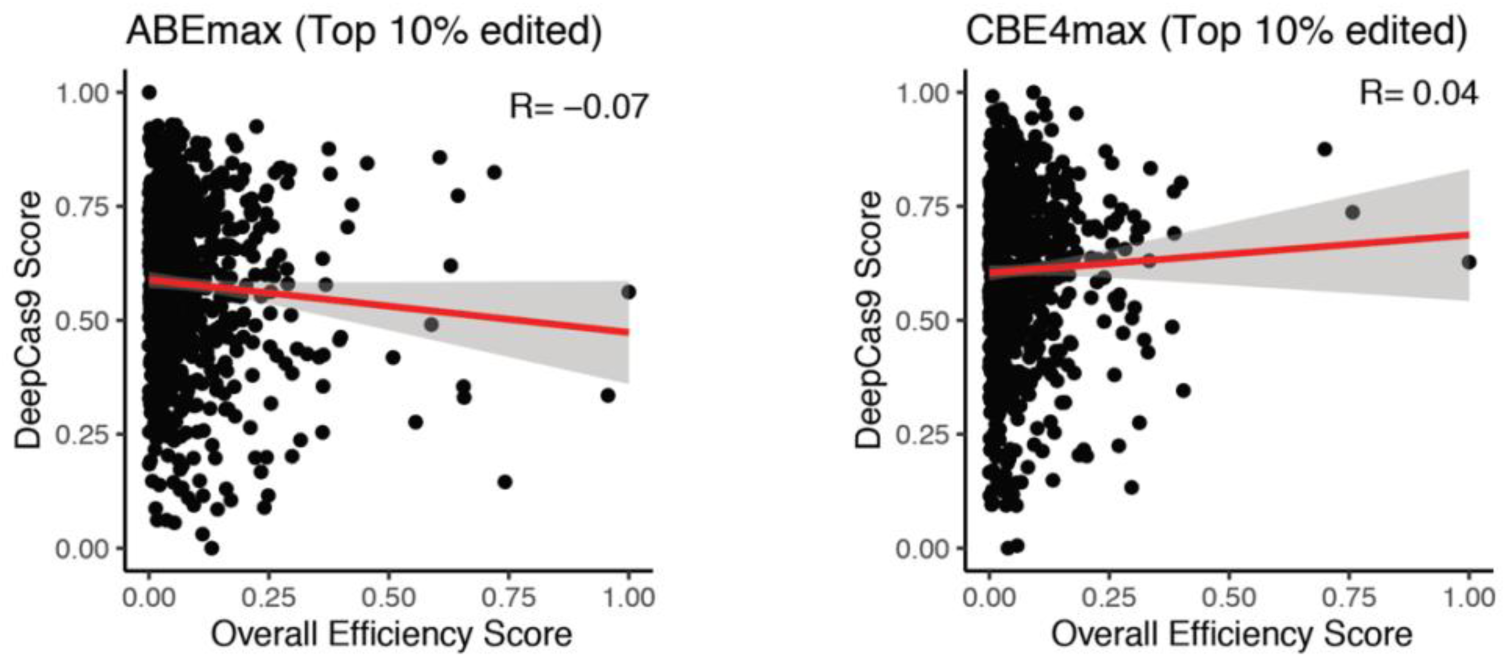
DeepSpCas9 score prediction versus base editing efficiency. Correlation between DeepSpCas9 score and the overall editing efficiency of top 10% edited sequences (ABEmax: n = 664; CBE4max: n = 901). The overall editing efficiency scores were scaled between 0 and 1.

**Supplemental figure 5:**
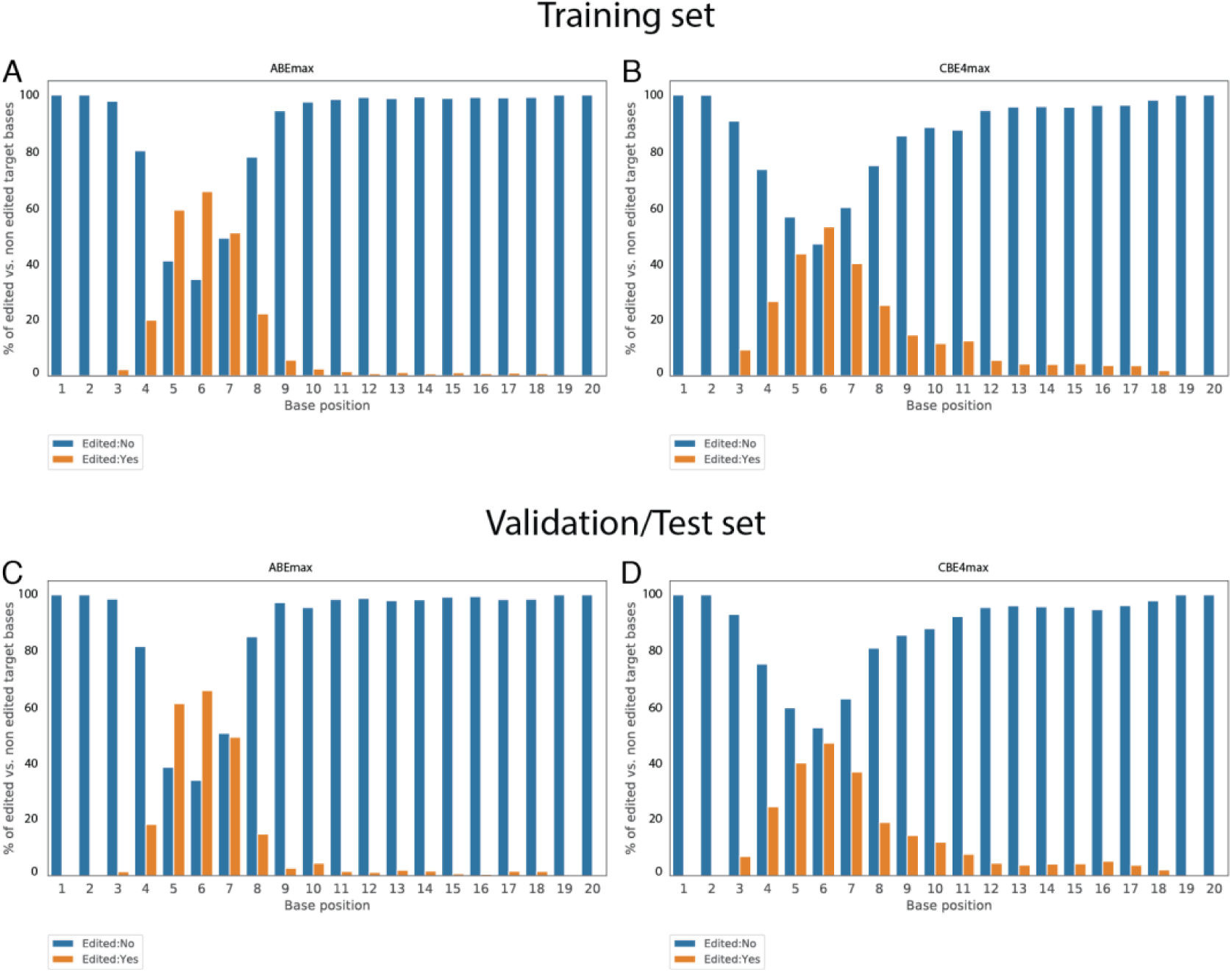
Per position base conversion (A-to-G; C-to-T) in the training (A, B) and validation/test (C, D) data set of the ABEmax and CBE4max screens.

**Supplemental figure 6:**
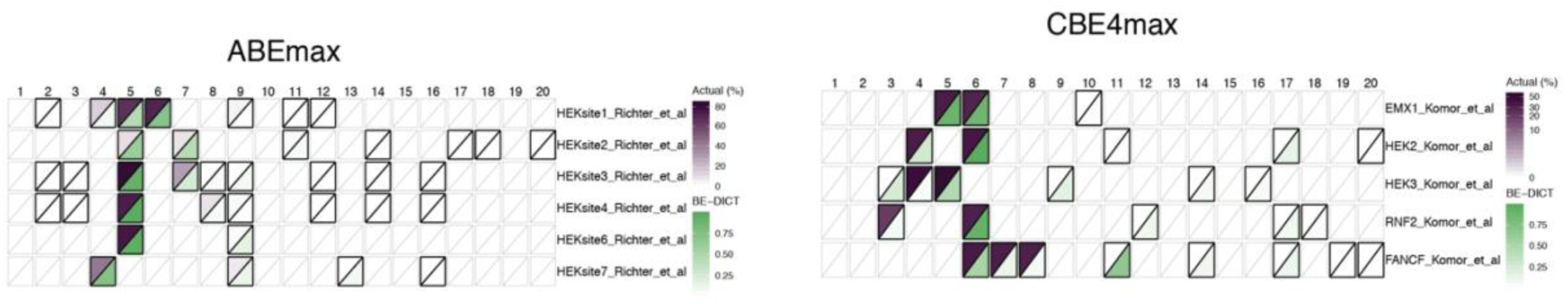
Comparison of predicted (BE-DICT) versus published base editing activities on genomic loci for ABEmax and CBE4max. Heatmap shows the BE-DICT prediction values (green) and the editing percentage (purple) for target sequences from Richter et al. [8] (ABEmax) and Komor et al. [19] (CBE4max).

**Supplemental figure 7:**
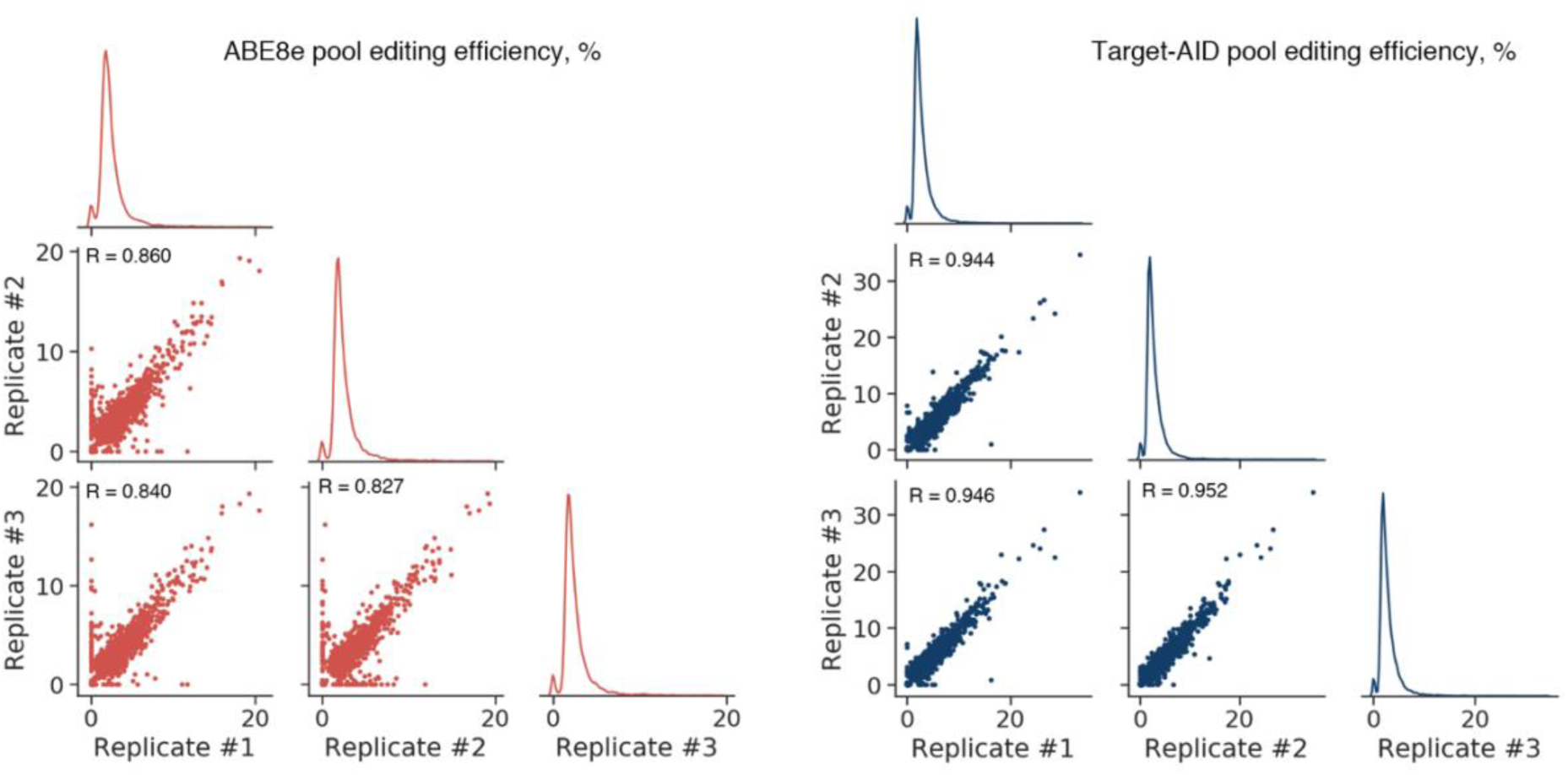
Correlations between the results of technical replicates of the high-throughput screening experiments. Scatter plots show the correlation between editing efficiencies for all sequences that were edited in at least one of the three technical replicates (with editing efficiency ≥ 1.5%) from experiments of library-cells treated with ABE8e and Target-AID.

**Supplemental figure 8:**
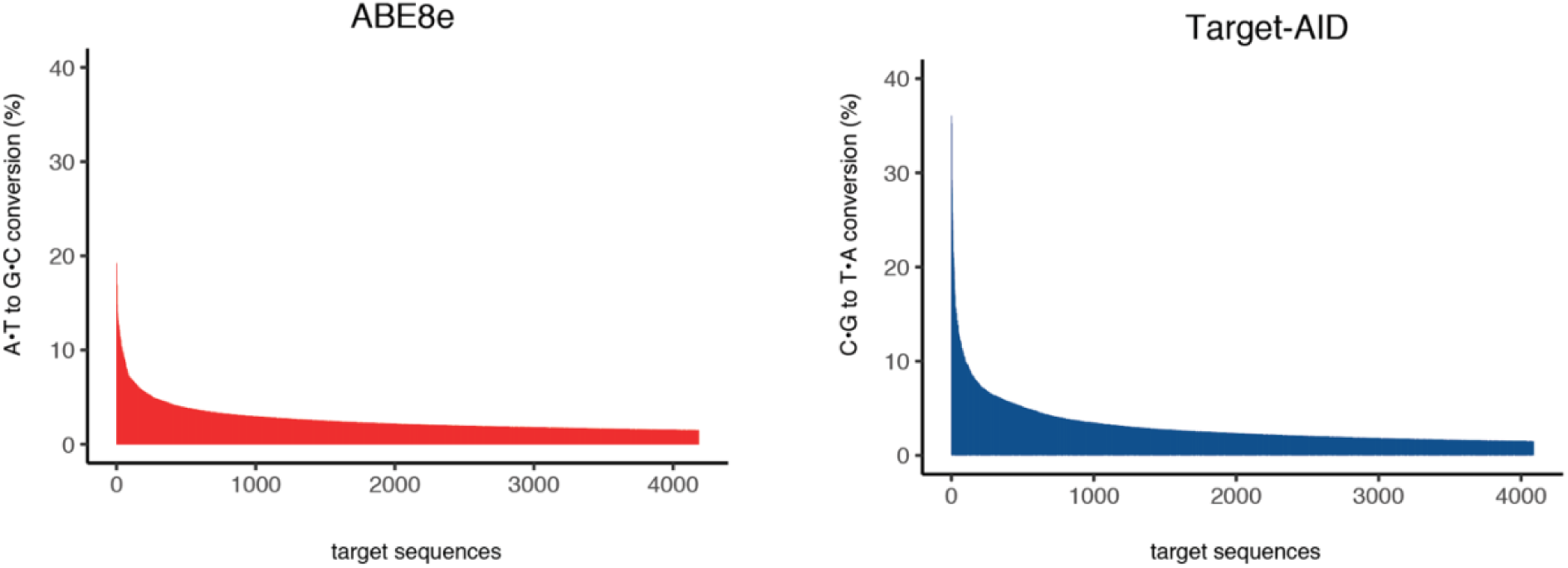
Editing rates in the self-targeting pooled library for ABE8e and Target-AID screens. Percentage of overall editing efficiency scores per target sequence for ABEmax and CBEmax.

**Supplemental figure 9:**
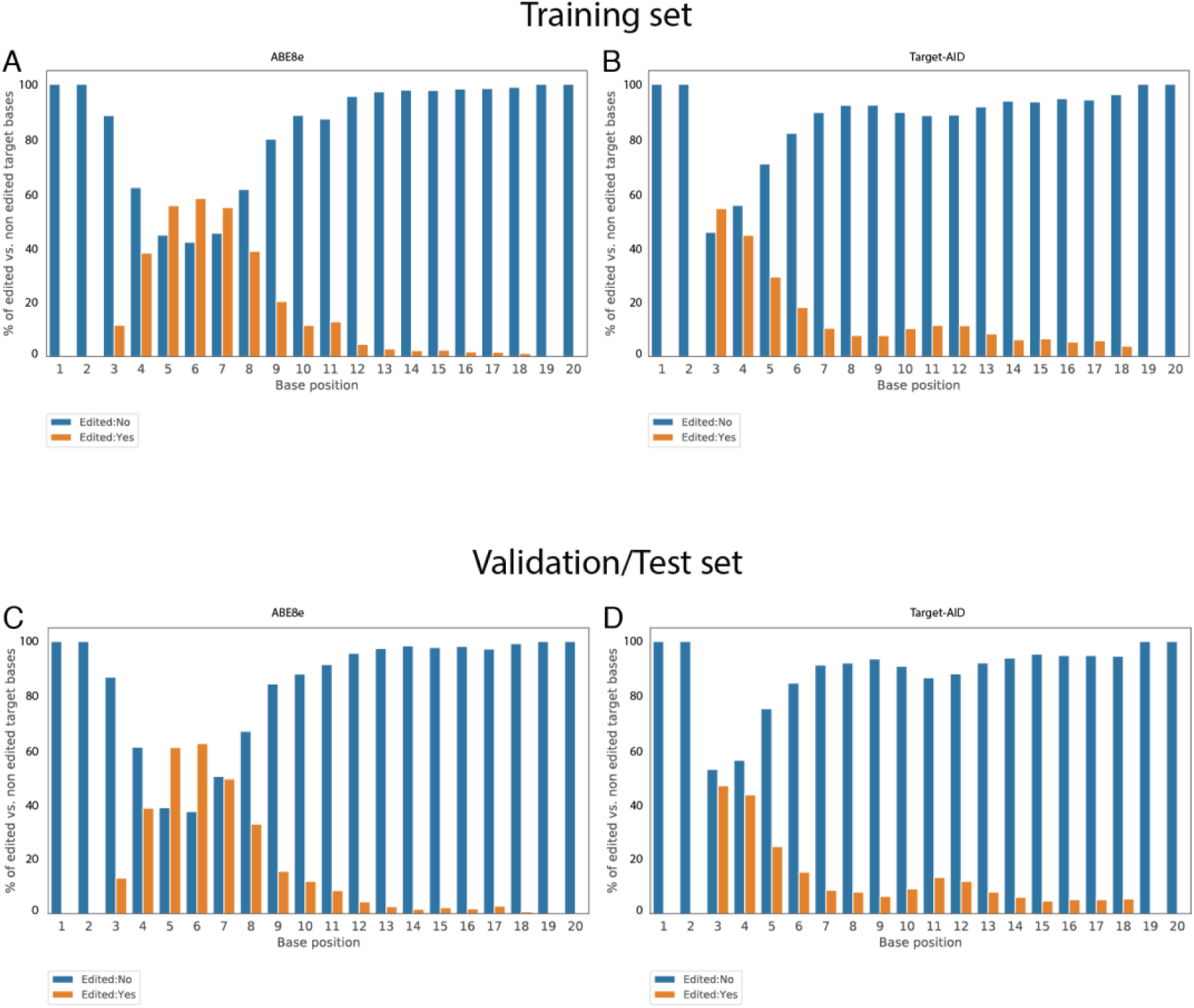
Per position base conversion (A-to-G; C-to-T) in the training (A, B) and validation/test (C, D) data set of the ABE8e and Target-AID screens.

**Supplemental figure 10:**
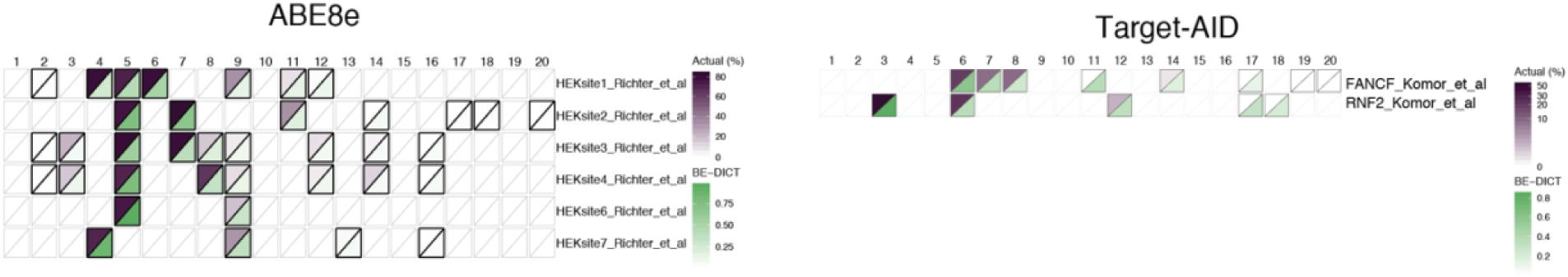
Comparison of predicted (BE-DICT) versus published base editing activities on genomic loci for ABE8e and Target-AID. Heatmap shows the BE-DICT prediction values (green) and the editing percentage (purple) for target sequences from Richter et al. [8] (ABE8e) and Komor et al. [19] (Target-AID).

**Supplemental figure 11:**
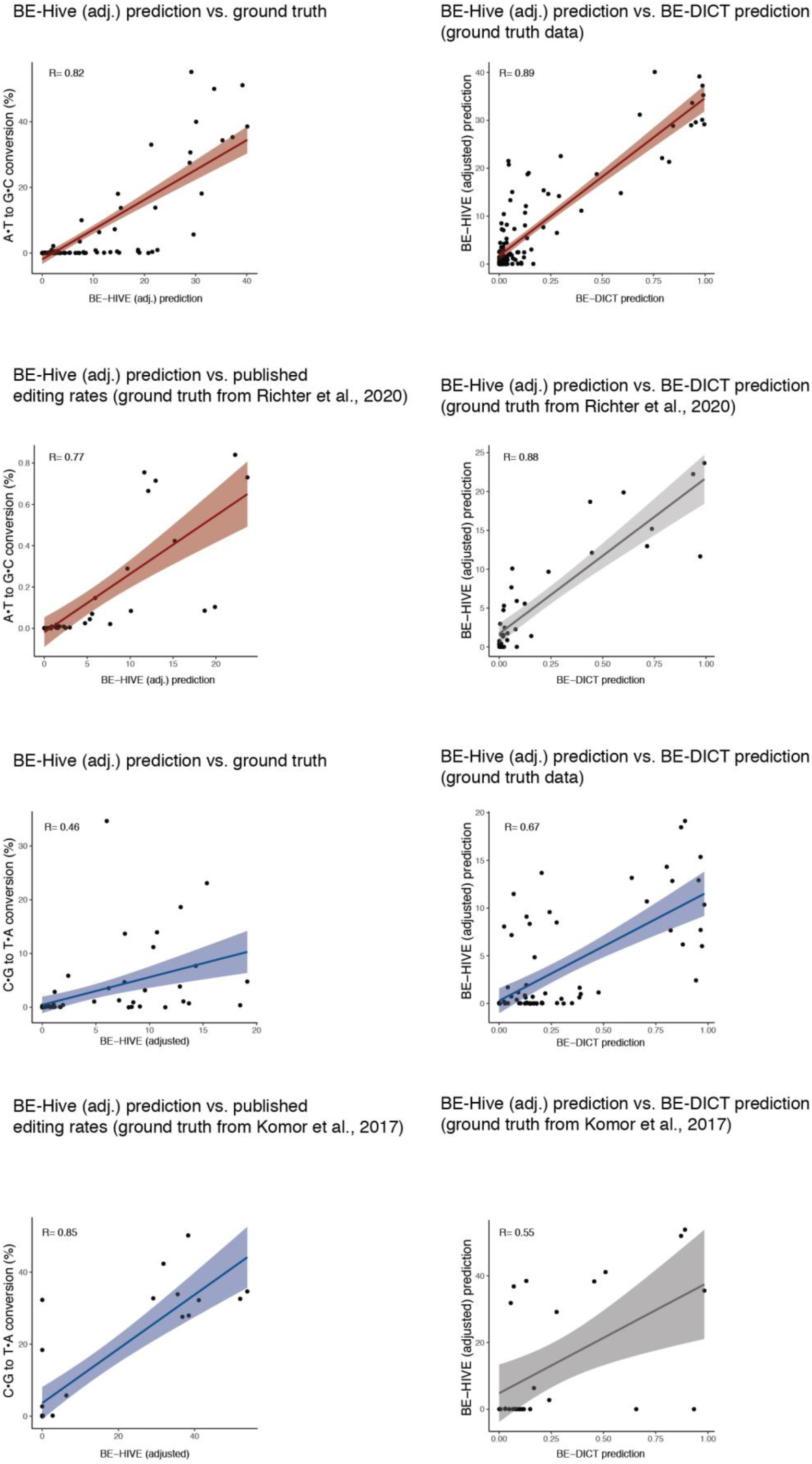
Comparison of BE-Hive and BE-DICT performance. Left panels: Correlation plots between the BE-Hive (adjusted) prediction score and editing percentages (A-to-G or C-toT). Same experimental data as shown in Fig.3b,c,e,f was plotted. Right panels: Correlation plot between the BE-Hive (adjusted) prediction score and the BE-DICT probability score. BE-Hive (adjusted) values were calculated as indicated in their online tool.

## Notes

### Competing Interest Statement

The authors have declared no competing interest.

