## Supplementary data file 1 for "Predicting base editing outcomes with an attention-based deep learning algorithm trained on high-throughput target library screens"

### 1 Attention-based Neural Network

#### 1.1 Model Overview

We designed and implemented a multi-head self-attention model (named BE-DICT) inspired by the Transformer [18] encoder architecture. BE-DICT is implemented in PyTorch [28] and takes a sequence of nucleotides (i.e. using protospacer sequence of 20 bp window) as input and computes the probability of editing for each target nucleotide as output. The target nucleotides in our experiments were base *A* for ABEmax and ABE8e editors, and *C* for CBE4max and Target-AID base editors respectively.
The model has three main blocks: An (1) **Embedding block** that embeds both the nucleotide's and its corresponding position from one-hot encoded representation to a dense vector representation (Panel A in Figure 1).
An (2) **Encoder block** that contains (a) a self-attention layer (with multi-head support), (b) layer normalization & residual connections (⇢), and (c) feed-forward network (Panel B in Figure 1).
Lastly, an (3) **Output block** that contains (a) a position attention layer and (b) a classifier layer (Panel C in Figure 1).
A formal description of each component of the model is described in their respective sections below.

#### 1.2 Embedding Block

Formally, given a protospacer sequence $\underline{S} = [x_1, x_2, \cdots, x_T]$, a nucleotide at position $t$ is represented by 1-of-*K* encoding where $K$ is the size of the set of all nucleotide letters in the data such that $x_t \in [0,1]^K$. An embedding matrix $W_e$ is used to map the input $x_t$ to a fixed-length vector representation (Eq. 1)

$$e_t = W_e x_t \tag{1}$$

where $W_e \in \mathbb{R}^{d_e \times K}$, $e_t \in \mathbb{R}^{d_e}$, and $d_e$ is the dimension of vector $e_t$.
Similarly, each nucleotide's position $p_t$ in the sequence $\underline{S}$ is represented by 1-of-*T* encoding where $T$ is the number of elements in the sequence (i.e. length of protospacer sequence) such that $p_t \in [0,1]^T$. An embedding matrix $W_{p'}$ is used to map the input $p_t$ to a fixed-length vector representation (Eq. 2)

$$p'_t = W_{p'} p_t \tag{2}$$

where $W_{p'} \in \mathbb{R}^{d_{p'} \times T}$, $p'_t \in \mathbb{R}^{d_{p'}}$ and $d_{p'}$ is the dimension of vector $p'_t$ such that $d_e$ and $d_{p'}$ were equal (denoted by $d$ from now on).
Both embeddings $e_t$ and $p'_t$ were summed (Eq. 3) to get a unified representation for every element in the sequence $\underline{S}$ (i.e. compute a new sequence $\underline{U} = [u_1, u_2, \cdots, u_T]$ where $u_t \in \mathbb{R}^d$, $\forall t \in [1, \cdots, T]$).

$$u_t = e_t + p'_t \tag{3}$$

#### 1.3 Encoder Block

##### 1.3.1 Self-Attention Layer

We followed a multi-head self-attention approach where multiple single-head self-attention layers are used in parallel (i.e. simultaneously) to process each

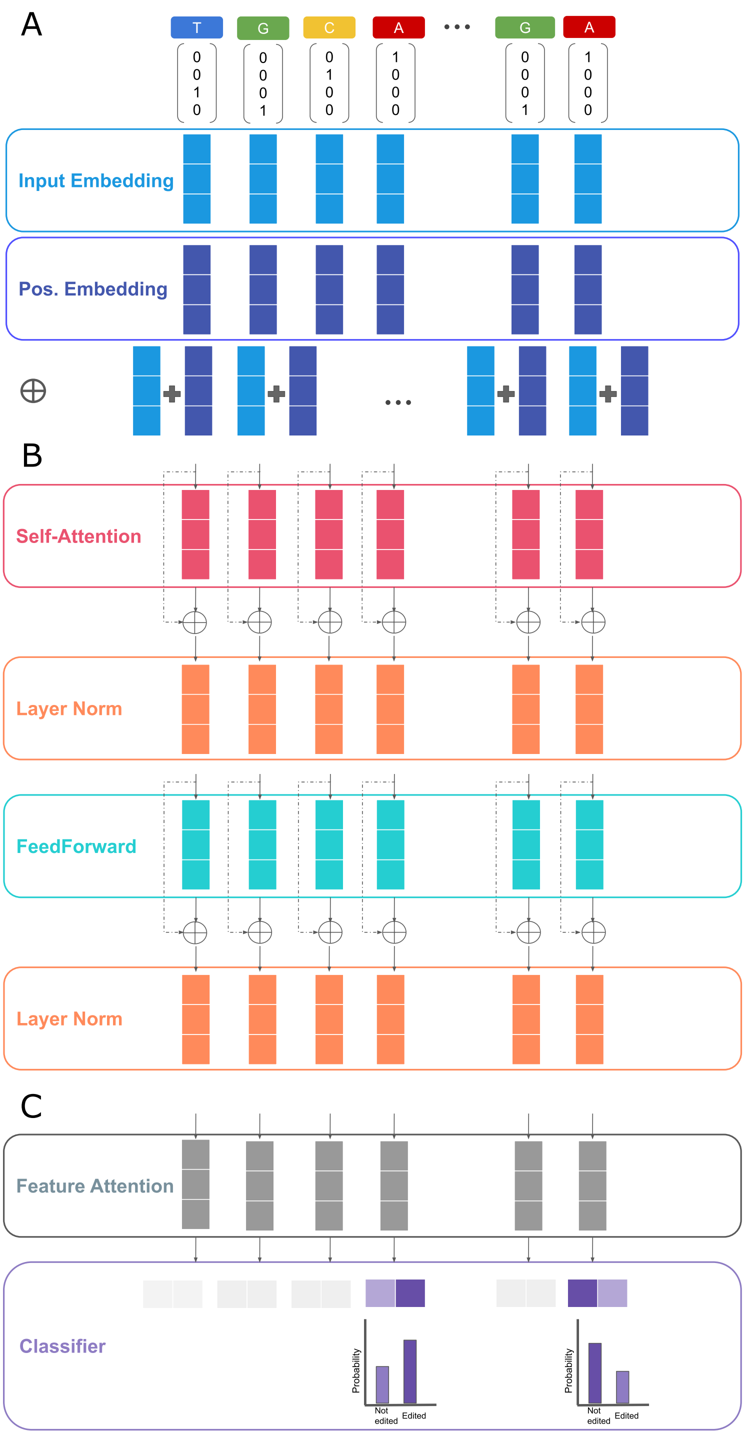


Figure 1: Overview of the BE-DICT attention-based neural network model.

input vector $u_t$. The outputs from every single-head layer are concatenated and transformed to generate a fixed-length vector using an affine transformation. The single-head self-attention approach [18] performs linear transformation to the input vector using three separate matrices: (1) a queries matrix $W_{query}$, (2) keys matrix $W_{key}$, and (3) values matrix $W_{value}$. Each input $u_t$ in $\underline{U}$ is mapped using these matrices to compute three new vectors (Eq. 4, 5, and 6)

$$q_t = W_{query} u_t \tag{4}$$

$$k_t = W_{key} u_t \tag{5}$$

$$v_t = W_{value} u_t \tag{6}$$

where $W_{query}$, $W_{key}$, $W_{value} \in \mathbb{R}^{d' \times d}$, $q_t$, $k_t$, $v_t \in \mathbb{R}^{d'}$ are query, key and value vectors, and $d'$ is the dimension of the three computed vectors respectively. In a second step, attention scores are computed using the pairwise similarity between the query and key vectors for each position $t$ in the sequence. The similarity is defined by computing a scaled dot-product between the pairwise vectors. At each position $t$, we compute attention scores $\alpha_{tl}$ representing the similarity between $q_t$ and vectors $k_l$ $\forall l \in [1, \dots, T]$ (Eq. 7, 8) normalized using $softmax$ function. Then a weighted sum using the attention scores $\alpha_{tl}$ and value vectors $v_l$ $\forall l \in [1, \dots, T]$ is performed (Eq. 9) to generate a new vector representation $z_t \in \mathbb{R}^{d'}$ at position $t$. This process is applied to every position in the sequence $\underline{U}$ to obtain a sequence of vectors $\underline{Z} = [z_1, z_2, \cdots, z_T]$.

$$\alpha_{tl} = \frac{\exp\left(score(q_t, k_l)\right)}{\sum_{l=1}^{T} \exp\left(score(q_t, k_l)\right)} \tag{7}$$

$$score(q_t, k_l) = \frac{{q_t}^{\top} k_l}{\sqrt{d'}} \tag{8}$$

$$z_t = \sum_{l=1}^{T} \alpha_{tl} v_l \tag{9}$$

In a multi-head setting with $H$ number of heads, the queries, keys and values matrices will be indexed by superscript $h$ (i.e. $W^h_{query}$, $W^h_{key}$, $W^h_{value} \in \mathbb{R}^{d' \times d}$) and applied separately to generate a new vector representation $z^h_t$ for every single-head self-attention layer. The output from each single-head layer is concatenated into one vector $z^{concat}_t = concat(z^1_t, z^2_t, \cdots, z^H_t)$ where $z^{concat}_t \in \mathbb{R}^{d'H}$ and then transformed using affine transformation (Eq. 10) such that $W_{unify} \in \mathbb{R}^{d' \times d'H}$ and $b_{unify} \in \mathbb{R}^{d'}$. This process is applied to each position in the sequence $\underline{Z}$ to generate a sequence $\underline{\tilde{Z}} = [\tilde{z}_1, \tilde{z}_2, \cdots, \tilde{z}_T]$.

$$\tilde{z}_t = W_{unify} z^{concat}_t + b_{unify} \tag{10}$$

###### 1.3.2 Layer Normalization & Residual Connections

We used residual connections / skip-connections (⇢) [27] in order to improve the gradient flow in layers during training. This is done by summing both the newly computed output of the current layer with the output from the previous layer. In our setting, a first residual connection sums the output of the self-attention layer $\tilde{z}_t$ and the output of embedding block $u_t$ for each position $t$ in

the sequence. We will refer to the summed output by $\tilde{z}_t$ for simplicity.
Layer normalization [26] was used in two occasions; (1) after the self-attention layer and (2) the feed-forward network layer with the goal to ameliorate the "covariate-shift" problem by re-standardizing the computed vector representations (i.e. using the mean and variance across the features/embedding dimension $d'$). Given a computed vector $\tilde{z}_t$, $LayerNorm$ function will standardize the input vector using the mean $\mu_t$ and variance $\sigma_t^2$ along the features dimension $d'$ and apply a scaling $\gamma$ and shifting step $\beta$ (Eq. 13). $\gamma$ and $\beta$ are learnable parameters and $\epsilon$ is small number added for numerical stability.

$$\mu_t = \frac{1}{d'} \sum_{j=1}^{d'} \tilde{z}_{tj} \tag{11}$$

$$\sigma_t^2 = \frac{1}{d'} \sum_{j=1}^{d'} (\tilde{z}_{tj} - \mu_t)^2 \tag{12}$$

$$LayerNorm(\tilde{z}_t) = \gamma \times \frac{\tilde{z}_t - \mu_t}{\sqrt{\sigma_t^2 + \epsilon}} + \beta \tag{13}$$

##### 1.3.3 FeedForward Layer

After a layer normalization step, a feed-forward network consisting of two affine transformation matrices and non-linear activation function is used to further compute/embed the learned vector representations from previous layers. The first transformation (Eq. 14) uses $W_{MLP1} \in \mathbb{R}^{\xi d' \times d'}$ and $b_{MLP1} \in \mathbb{R}^{\xi d'}$ to transform input $\tilde{z}_t$ to new vector $\in \mathbb{R}^{\xi d'}$ where $\xi \in \mathbb{N}$ is multiplicative factor. A non-linear function such as $ReLU(z) = max(0, z)$ is applied followed by another affine transformation using $W_{MLP2} \in \mathbb{R}^{d' \times \xi d'}$ and $b_{MLP2} \in R^{d'}$ to obtain vector $r_t \in \mathbb{R}^{d'}$. A layer normalization (Eq. 15) is applied to obtain $\tilde{r}_t \in \mathbb{R}^{d'}$.

$$r_t = W_{MLP2} ReLU(W_{MLP1} \tilde{z}_t + b_{MLP1}) + b_{MLP2} \tag{14}$$

$$\tilde{r}_t = LayerNorm(r_t) \tag{15}$$

These transformations are applied to each vector in sequence $\underline{\tilde{Z}}$ to obtain new sequence $\underline{R} = [\tilde{r}_1, \tilde{r}_2, \cdots, \tilde{r}_T]$. At this point, the *encoder* block operations are done and multiple encoder blocks can be stacked in series for $E$ number of times. In our experiments, $E$ was a hyperparameter that was empirically determined using a validation set (as the case of the number of attention heads $H$ used in self-attention layer).

#### 1.4 Output Block

##### 1.4.1 Position Attention Layer

The position attention layer is parametrized by a set of *global* context vectors $\underline{C} = [c_1, c_2, \cdots, c_T]$ corresponding to each position in the protospacer sequence. These context vectors are learnable parameters optimized during the training. For a target base at position $t$, attention scores $\psi_t \forall t \in [1, \cdots, T]$ are calculated using the pairwise similarity between the context vector $c_t \in \mathbb{R}^{d'}$ with the vectors $[\tilde{r}_1, \tilde{r}_2, \cdots, \tilde{r}_T]$ computed in the previous layer and representing every position in the protospacer sequence (Eq. 16, 17). These scores are normalized and used to compute a weighted sum of the $[\tilde{r}_1, \tilde{r}_2, \cdots, \tilde{r}_T]$ vectors to generate a new vector representation $o_t \in \mathbb{R}^{d'}$ at position $t$. This process is done at each target position to generate a sequence $\underline{O} = [o_1, o_2, \cdots, o_T]$ that is further passed to the classifier layer.

$$\psi_t = \frac{\exp(score(c_t, \tilde{r}_t))}{\sum_{j=1}^{T} \exp(score(c_t, \tilde{r}_j))} \tag{16}$$

$$score(c_t, \tilde{r}_t) = \frac{{c_t}^{\top} \tilde{r}_t}{\sqrt{d'}} \tag{17}$$

$$o_t = \sum_{t=1}^{T} \psi_t \tilde{r}_t \tag{18}$$

###### 1.4.2 Output Classifier

The last layer in the model takes as input the computed representation vectors $\underline{O} = [o_1, o_2, \cdots, o_T]$ from the *position attention* layer and performs an affine transformation followed by *softmax* operation to compute a probability distribution on the outcomes (i.e. edit vs. no edit) for every target base under consideration (i.e. base *A* or *C*). That is, the probability distribution $\hat{y}_t$ at position $t$ is computed using Eq. 19

$$\hat{y}_t = \sigma(W_o o_t + b_o) \tag{19}$$

where $W_o \in \mathbb{R}^{|V_{outcome}| \times d'}$, $b_o \in \mathbb{R}^{|V_{outcome}|}$, $V_{outcome} \in \{0, 1\}$ is the set of admissible labels (binary variable in our case). Moreover, $|V_{outcome}|$ is the number of labels, $d'$ is the dimension of $o_t$ and $\sigma$ is the $softmax$ function.

##### 1.5 Objective Function

We defined the loss for an $i$-th sequence at each position $t$ by the cross-entropy loss

$$l_t^{(i)} = -\sum_{c=1}^{|V_{outcome}|} y_{t,c}^{(i)} \times log(\hat{y}_{t,c}^{(i)}) \tag{20}$$

where the loss for the $i$-th sequence is defined by the average loss over the sequence length $T$

$$L_i = \frac{1}{T} \sum_{t=1}^{T} l_t^{(i)} \tag{21}$$

Given that our focus is on target nucleotides (i.e. bases *A* or *C* depending on the base editor used), our model's focus should be on positions where target bases occur. Hence, we modified the defined loss over $i$-th sequence (see Eq. 21) by defining an average loss for target base $L_i^{target}$ (Eq. 22)

$$L_i^{target} = \frac{1}{\sum_{t=1}^{T} \mathbb{1}[x_{t,base}^{(i)} = 1]} \sum_{t=1}^{T} l_t^{(i)} \mathbb{1}[x_{t,base}^{(i)} = 1] \tag{22}$$

where $\mathbb{1}[x_{t,base}^{(i)} = 1]$ is an indicator function that is equal to $1$ when $x_t^i$ representing the nucleotide at position $t$ for the $i$-th sequence is the target base (i.e. *A* for ABEmax and ABE8e and *C* for CBE4max and Target-AID base editors respectively). Lastly, the objective function for the whole training set $D_{train}$ is defined by the average loss across all the sequences in $D_{train}$ plus a weight regularization term (i.e. $l_2$-norm regularization) applied to the model parameters represented by $\boldsymbol{\theta}$

$$L(\boldsymbol{\theta}) = \frac{1}{N}\sum_{i=1}^{N} L_i^{target} + \frac{\lambda}{2}||\boldsymbol{\theta}||_2^2 \tag{23}$$

The training is done using mini-batches where computing the loss function and updating the parameters/weight occur after processing each mini-batch of the training set.

### 2 Experiments

#### 2.1 Training & Evaluation Workflow

For model training we used ≈ 80% of the dataset consisting of randomized-DNA target sequences, and performed stratified random splits for the rest of the sequences to generate an equal ratio (1:1) between test and validation datasets. We repeated this process five times (denoted by runs), in which we trained and evaluated a model for each base editor separately for each run. Due to the imbalance in outcome classes, training examples were weighted inversely proportional to class/outcome frequencies in the training data. BE-DICT performance was evaluated using area under the receiver operating characteristic curve (AUC), and area under the precision recall curve (AUPR). During training of the models, the epoch in which the model achieved the best AUPR on the validation set was recorded, and model state as it was trained up to that epoch was saved. This best model, as determined by the validation set, was then tested on the test split. The evaluation of the trained models for each base editor was based on their average performance on the test sets across the five runs.
We further compared BE-DICT's prediction accuracy at each position with a majority class predictor (one for each base editor) that uses the training data to estimate the per position prior (i.e. base rate) probability of having a target base edited. Based on the estimated prior, the predictor assigns the majority class (one with higher probability) as the outcome for the target base at the position under consideration.

#### 2.2 Hyperparameters Optimization

We used a uniform random search strategy [29] that randomly chose a set of hyperparameters configurations (i.e. embedding dimension, number of attention heads, dropout probability, etc.) from the set of all possible configurations. Then the best configuration for each model (i.e. the one achieving best performance on the validation set) was used for the final training and testing.
The range of possible hyperparameters configuration (i.e. choice of values for

hyperparameters) for BE-DICT models is reported in Table 1. In our experiments, there was an overlap between the best hyperparameter configurations for the different models trained on the various base editors. Hence, we opted for a common hyperparameter configuration among all models.

**List 1** BE-DICT hyperparameters options

- Embedding Block operations
  - Nucleotide embedding layer
    - embedding dimension $d$ ........................ $\{16, 32, 64, 128\}$
  - Position embedding layer
    - embedding dimension $d$ ........................ $\{16, 32, 64, 128\}$
- Encoder Block operations
  - Self-attention layer
    - embedding dimension $d'$ ........................ $\{16, 32, 64, 128\}$
    - Number of attention heads $H$ .................. $\{4, 6, 8, 12, 16\}$
    - Dropout .......................................... $\{0.1, 0.3, 0.5\}$
  - Feed-Forward layer
    - MLP embedding factor (multiplier) $\xi$ ................. $\{2\}$
    - Non-linear function ................... $\{tanh, ELU, ReLU\}$
  - Number of repeats for Encoder Block $E$ ................ $\{2, 4, 6\}$
- $l_2$-norm regularization $\lambda$ ................ $\{10^{-5}, 10^{-4}, 10^{-3}, 10^{-2}, 10^{-1}\}$
- Batch size during training $|B|$ ............. $\{1000, 2000, 3000, 4000\}$
- Optimization algorithm .................................. $\{Adam\}$

### 3 DNA used in this study

#### 3.1 Oligonucleotide pool design

Oligonucleotide sequences, each containing a guide RNA and corresponding target sequence pair, used to generate the pooled library.

Primer binding sites
Guide RNA sequence
Spcas9 guide RNA scaffold
Target sequence with PAM
Randomized barcode 1 and randomized barcode 2

ATCTTGTGGAAAGGACGAAACACCGNNNNNNNNNNNNNNNNNNNNGTTTTAGA
GCTAGAAATAGCAAGTTAAAATAAGGCTAGTCCGTTATCAACTTGAAAAAGTGG
CACCGAGTCGGTGCTTTTTTNNNNNNNNNNNNNNNNNNNNNNNNNNNGGNNN
NNNGCTCTACCACTTGTACTTCAGC

#### 3.2 Primers used for high-throughput sequencing

| **Primer used for:** | | **Sequence (5'-3')** |
|---|---|---|
| Oligo-pool amplification | FW | ATCTTGGGAAAGGACGAAACACC |
| | RV | GCTGAAGTACAAGTGGTAGAGC |
| Pooled screens (PCR on genomic DNA) | FW | CTTTCCCTACACGACGCTCTTCCGATCTNNNNNNNNGAAAAAGTGGCACCGAGTCG |
| | RV | GGAGTTCAGACGTGTGCTCTTCCGATCTNNNNNNNNACTATTCTTTCCCCTGCACTGT |
| DOCK3 no2 | FW | CTTTCCCTACACGACGCTCTTCCGATCTNNNNTCCTACATTTCAGTGAGCGGT |
| | RV | GGAGTTCAGACGTGTGCTCTTCCGATCTNNNNGCAGGCCACTGGTTAGAGTC |
| TARDBP no2 | FW | CTTTCCCTACACGACGCTCTTCCGATCTNNNNGCGCTGTACAGAGGACATGA |
| | RV | GGAGTTCAGACGTGTGCTCTTCCGATCTNNNNCCTGAATGGCTTGGGGATGA |
| ZNF212 no1 | FW | CTTTCCCTACACGACGCTCTTCCGATCTNNNNGCCTGCACAGTGAGGAAGAG |
| | RV | GGAGTTCAGACGTGTGCTCTTCCGATCTNNNNCCAGAGCCTGTTGGAAGCC |

| | | |
|---|---|---|
| NEK1 no2 | FW | CTTTCCCTACACGACGCTCTTCCGATCTNNNNAGCCATGCTTTTGATGTACGT |
| | RV | GGAGTTCAGACGTGTGCTCTTCCGATCTNNNNAACAGATCCCCTCCCTCACA |
| RSF1 no1 | FW | CTTTCCCTACACGACGCTCTTCCGATCTNNNNCCCCTTCTCTCCTCCCCTTC |
| | RV | GGAGTTCAGACGTGTGCTCTTCCGATCTNNNNTTGTTCTCGAGAAGTCCCGC |
| RSF1 no2 | FW | CTTTCCCTACACGACGCTCTTCCGATCTNNNNAGCTAGAAAAACCTTTGCCAGA |
| | RV | GGAGTTCAGACGTGTGCTCTTCCGATCTNNNNGGCTTCTCCATGCTACTTTTGG |
| FANCF no3 | FW | CTTTCCCTACACGACGCTCTTCCGATCTNNNNCTACCTGCGCCACATCCATC |
| | RV | GGAGTTCAGACGTGTGCTCTTCCGATCTNNNNTTCGCTAATCCCGGAACTGG |
| FANCF no5 | FW | CTTTCCCTACACGACGCTCTTCCGATCTNNNNCTACCTACGTCAGCACCTGG |
| | RV | GGAGTTCAGACGTGTGCTCTTCCGATCTNNNNGATGGATGTGGCGCAGGTAG |
| EMX1 no1 | FW | CTTTCCCTACACGACGCTCTTCCGATCTNNNNCAGGTGAAGGTGTGGTTCCA |
| | RV | GGAGTTCAGACGTGTGCTCTTCCGATCTNNNNCACCGGTTGATGTGATGGGA |
| EMX1 no2 | FW | CTTTCCCTACACGACGCTCTTCCGATCTNNNNATAGTCCCCTTGGGGTGACA |
| | RV | GGAGTTCAGACGTGTGCTCTTCCGATCTNNNNCCGGCCAGAGACTTCCTGTA |
| HEK site no1 | FW | CTTTCCCTACACGACGCTCTTCCGATCTNNNNCCAGCCCCATCTGTCAAACT |
| | RV | GGAGTTCAGACGTGTGCTCTTCCGATCTNNNNTGAATGGATTCCTTGGAAACAATGA |
| HEK site no2 | FW | CTTTCCCTACACGACGCTCTTCCGATCTNNNNAGAGACTGATTGCGTGGAGT |
| | RV | GGAGTTCAGACGTGTGCTCTTCCGATCTNNNNCACTCCAGCCTAGGCAACAA |
| HEK site no3 | FW | CTTTCCCTACACGACGCTCTTCCGATCTNNNNCCGACAGCCAGTGGTTAAGT |
| | RV | GGAGTTCAGACGTGTGCTCTTCCGATCTNNNNGCTTTTCACCGACTGCACAG |
| HEK site no8 | FW | CTTTCCCTACACGACGCTCTTCCGATCTNNNNCCCTGTTCCTAAAGCCCACC |
| | RV | GGAGTTCAGACGTGTGCTCTTCCGATCTNNNNACTGGTTCTGTTTGTGGCCA |
| HEK site no9 | FW | CTTTCCCTACACGACGCTCTTCCGATCTNNNNTTGCTTATTGCTGAGGGGCA |

| | | |
|---|---|---|
| | RV | GGAGTTCAGACGTGTGCTCTTCCGATCTNNNNACCTCTCTCCTCCAGCTGAG |
| HEK site no10 | FW | CTTTCCCTACACGACGCTCTTCCGATCTNNNNTCCACCTCCCCACTTCTCTT |
| | RV | GGAGTTCAGACGTGTGCTCTTCCGATCTNNNNGGTGAAATGAGCAAGGCACA |
| HEK site no11 | FW | CTTTCCCTACACGACGCTCTTCCGATCTNNNNCCCTAAACCACCTGCAGAGG |
| | RV | GGAGTTCAGACGTGTGCTCTTCCGATCTNNNNCAGCCCCAGCCACATTCTAT |
| HEK site no14 | FW | CTTTCCCTACACGACGCTCTTCCGATCTNNNNGAACCTGAAGCCTTTCCCCA |
| | RV | GGAGTTCAGACGTGTGCTCTTCCGATCTNNNNAACCTGTGTGACACTTGGCA |
| HEK site no16 | FW | CTTTCCCTACACGACGCTCTTCCGATCTNNNNGGGAGGTGGAGAGAGGATGT |
| | RV | GGAGTTCAGACGTGTGCTCTTCCGATCTNNNNTCCTGAGGTCTAGGAACCCG |
| HEK site no18 | FW | CTTTCCCTACACGACGCTCTTCCGATCTNNNNGCATTACCTGGGAGCCTGTT |
| | RV | GGAGTTCAGACGTGTGCTCTTCCGATCTNNNNAACTTCAGCGGGCATCAGAA |

#### 3.3 Oligonucleotides used for sgRNA cloning

| **Oligonucleotide used for cloning of sgRNA:** | | **Sequence (5'-3')** |
|---|---|---|
| DOCK3 no2 | FW | caccgTAAGACTGAACAAGAATGGT |
| | RV | aaacACCATTCTTGTTCAGTCTTAC |
| TARDBP no2 | FW | caccgCGGGAGTTCTTCTCTCAGTA |
| | RV | aaacTACTGAGAGAAGAACTCCCGC |
| ZNF212 no1 | FW | caccgTGCACCTGGCATCAACAcgg |
| | RV | aaacCCGTGTTGATGCCAGGTGCAC |
| NEK1 no2 | FW | caccgTGGCTCTCTCTACATAGTAA |
| | RV | aaacTTACTATGTAGAGAGAGCCAC |
| RSF1 no1 | FW | caccgTTCATTCCCCCTGTCACACG |
| | RV | aaacCGTGTGACAGGGGGAATGAAC |
| RSF1 no2 | FW | caccgACCCATTAAAGTTGAGGTGA |
| | RV | aaacTCACCTCAACTTTAATGGGTC |
| FANCF no3 | FW | caccgAGCGGCGGCTGCACAACCAG |
| | RV | aaacCTGGTTGTGCAGCCGCCGCTC |
| FANCF no5 | FW | caccgAGGCCCGGCGCACGGTGGCG |
| | RV | aaacCGCCACCGTGCGCCGGGCCTC |
| EMX1 no1 | FW | caccgGAGTCCGAGCAGAAGAAGAA |
| | RV | aaacTTCTTCTTCTGCTCGGACTCC |

| | | |
|---|---|---|
| EMX1 no2 | FW | caccgAGATTTATGCAAACGGGTTG |
| | RV | aaacCAACCCGTTTGCATAAATCTC |
| HEK site no1 | FW | caccgGAACACAAAGCATAGACTGC |
| | RV | aaacGCAGTCTATGCTTTGTGTTCC |
| HEK site no2 | FW | caccgGAGTATGAGGCATAGACTGC |
| | RV | aaacGCAGTCTATGCCTCATACTCC |
| HEK site no3 | FW | caccgGTCAAGAAAGCAGAGACTGC |
| | RV | aaacGCAGTCTCTGCTTTCTTGACC |
| HEK site no8 | FW | caccgGTAAACAAAGCATAGACTGA |
| | RV | aaacTCAGTCTATGCTTTGTTTACC |
| HEK site no9 | FW | caccgGAAGACCAAGGATAGACTGC |
| | RV | aaacGCAGTCTATCCTTGGTCTTCC |
| HEK site no10 | FW | caccgGAACATAAAGAATAGAATGA |
| | RV | aaacTCATTCTATTCTTTATGTTCC |
| HEK site no11 | FW | caccgGGACAGGCAGCATAGACTGT |
| | RV | aaacACAGTCTATGCTGCCTGTCCC |
| HEK site no14 | FW | caccgGGCTAAAGACCATAGACTGT |
| | RV | aaacACAGTCTATGGTCTTTAGCCC |
| HEK site no16 | FW | caccgGGGAATAAATCATAGAATCC |
| | RV | aaacGGATTCTATGATTTATTCCCC |
| HEK site no18 | FW | caccgACACACACACTTAGAATCTG |
| | RV | aaacCAGATTCTAAGTGTGTGTGTC |

#### 3.4 Sequences of SpCas9-base-editor plasmids used in this study

(a) CMV-ABEmax-P2A-GFP (Addgene no.112101)

NLS
Linker
TadA-TadA*
Nickase SpCas9(D10A)
P2A-GFP

ATGAAACGGACAGCCGACGGAAGCGAGTTCGAGTCACCAAAGAAGAAGCGGA
AAGTC_TCTGAAGTCGAGTTTAGCCACGAGTATTGGATGAGGCACGCACTGACC
CTGGCAAAGCGAGCATGGGATGAAAGAGAAGTCCCCGTGGGCGCCGTGCTGG
TGCACAACAATAGAGTGATCGGAGAGGGATGGAACAGGCCAATCGGCCGCCA
CGACCCTACCGCACACGCAGAGATCATGGCACTGAGGCAGGGAGGCCTGGTC
ATGCAGAATTACCGCCTGATCGATGCCACCCTGTATGTGACACTGGAGCCATG
CGTGATGTGCGCAGGAGCAATGATCCACAGCAGGATCGGAAGAGTGGTGTTCG
GAGCACGGGACGCCAAGACCGGCGCAGCAGGCTCCCTGATGGATGTGCTGCA
CCACCCCGGCATGAACCACCGGGTGGAGATCACAGAGGGAATCCTGGCAGAC

GAGTGCGCCGCCCTGCTGAGCGATTTCTTTAGAATGCGGAGACAGGAGATCAA
GGCCCAGAAGAAGGCACAGAGCTCCACCGACTCTGGAGGATCTAGCGGAGGA
TCCTCTGGAAGCGAGACACCAGGCACAAGCGAGTCCGCCACACCAGAGAGCT
CCGGCGGCTCCTCCGGAGGATCCTCTGAGGTGGAGTTTTCCCACGAGTACTGG
ATGAGACATGCCCTGACCCTGGCCAAGAGGGCACGCGATGAGAGGGAGGTGC
CTGTGGGAGCCGTGCTGGTGCTGAACAATAGAGTGATCGGCGAGGGCTGGAA
CAGAGCCATCGGCCTGCACGACCCAACAGCCCATGCCGAAATTATGGCCCTGA
GACAGGGCGGCCTGGTCATGCAGAACTACAGACTGATTGACGCCACCCTGTAC
GTGACATTCGAGCCTTGCGTGATGTGCGCCGGCGCCATGATCCACTCTAGGAT
CGGCCGCGTGGTGTTTGGCGTGAGGAACGCAAAAACCGGCGCCGCAGGCTCC
CTGATGGACGTGCTGCACTACCCCGGCATGAATCACCGCGTCGAAATTACCGA
GGGAATCCTGGCAGATGAATGTGCCGCCCTGCTGTGCTATTTCTTTCGGATGC
CTAGACAGGTGTTCAATGCTCAGAAGAAGGCCCAGAGCTCCACCGAC_TCCGG
AGGATCTAGCGGAGGCTCCTCTGGCTCTGAGACACCTGGCACAAGCGAGAGC
GCAACACCTGAAAGCAGCGGGGGCAGCAGCGGGGGGTCA_GACAAGAAGTAC
AGCATCGGCCTGGCCATCGGCACCAACTCTGTGGGCTGGGCCGTGATCACCG
ACGAGTACAAGGTGCCCAGCAAGAAATTCAAGGTGCTGGGCAACACCGACCGG
CACAGCATCAAGAAGAACCTGATCGGAGCCCTGCTGTTCGACAGCGGCGAAAC
AGCCGAGGCCACCCGGCTGAAGAGAACCGCCAGAAGAAGATACACCAGACGG
AAGAACCGGATCTGCTATCTGCAAGAGATCTTCAGCAACGAGATGGCCAAGGT
GGACGACAGCTTCTTCCACAGACTGGAAGAGTCCTTCCTGGTGGAAGAGGATA
AGAAGCACGAGCGGCACCCCATCTTCGGCAACATCGTGGACGAGGTGGCCTA
CCACGAGAAGTACCCCACCATCTACCACCTGAGAAAGAAACTGGTGGACAGCA
CCGACAAGGCCGACCTGCGGCTGATCTATCTGGCCCTGGCCCACATGATCAAG
TTCCGGGGCCACTTCCTGATCGAGGGCGACCTGAACCCCGACAACAGCGACG
TGGACAAGCTGTTCATCCAGCTGGTGCAGACCTACAACCAGCTGTTCGAGGAA
AACCCCATCAACGCCAGCGGCGTGGACGCCAAGGCCATCCTGTCTGCCAGACT
GAGCAAGAGCAGACGGCTGGAAAATCTGATCGCCCAGCTGCCCGGCGAGAAG
AAGAATGGCCTGTTCGGAAACCTGATTGCCCTGAGCCTGGGCCTGACCCCCAA
CTTCAAGAGCAACTTCGACCTGGCCGAGGATGCCAAACTGCAGCTGAGCAAGG
ACACCTACGACGACGACCTGGACAACCTGCTGGCCCAGATCGGCGACCAGTAC
GCCGACCTGTTTCTGGCCGCCAAGAACCTGTCCGACGCCATCCTGCTGAGCGA
CATCCTGAGAGTGAACACCGAGATCACCAAGGCCCCCCTGAGCGCCTCTATGA
TCAAGAGATACGACGAGCACCACCAGGACCTGACCCTGCTGAAAGCTCTCGTG
CGGCAGCAGCTGCCTGAGAAGTACAAAGAGATTTTCTTCGACCAGAGCAAGAA
CGGCTACGCCGGCTACATTGACGGCGGAGCCAGCCAGGAAGAGTTCTACAAG
TTCATCAAGCCCATCCTGGAAAAGATGGACGGCACCGAGGAACTGCTCGTGAA
GCTGAACAGAGAGGACCTGCTGCGGAAGCAGCGGACCTTCGACAACGGCAGC
ATCCCCCACCAGATCCACCTGGGAGAGCTGCACGCCATTCTGCGGCGGCAGG
AAGATTTTTACCCATTCCTGAAGGACAACCGGGAAAAGATCGAGAAGATCCTGA
CCTTCCGCATCCCCTACTACGTGGGCCCTCTGGCCAGGGGAAACAGCAGATTC
GCCTGGATGACCAGAAAGAGCGAGGAAACCATCACCCCCTGGAACTTCGAGGA
AGTGGTGGACAAGGGCGCTTCCGCCCAGAGCTTCATCGAGCGGATGACCAACT
TCGATAAGAACCTGCCCAACGAGAAGGTGCTGCCCAAGCACAGCCTGCTGTAC
GAGTACTTCACCGTGTATAACGAGCTGACCAAAGTGAAATACGTGACCGAGGG
AATGAGAAAGCCCGCCTTCCTGAGCGGCGAGCAGAAAAAGGCCATCGTGGAC
CTGCTGTTCAAGACCAACCGGAAAGTGACCGTGAAGCAGCTGAAAGAGGACTA
CTTCAAGAAAATCGAGTGCTTCGACTCCGTGGAAATCTCCGGCGTGGAAGATC

GGTTCAACGCCTCCCTGGGCACATACCACGATCTGCTGAAAATTATCAAGGACA
AGGACTTCCTGGACAATGAGGAAAACGAGGACATTCTGGAAGATATCGTGCTG
ACCCTGACACTGTTTGAGGACAGAGAGATGATCGAGGAACGGCTGAAAACCTA
TGCCCACCTGTTCGACGACAAAGTGATGAAGCAGCTGAAGCGGCGGAGATACA
CCGGCTGGGGCAGGCTGAGCCGGAAGCTGATCAACGGCATCCGGGACAAGCA
GTCCGGCAAGACAATCCTGGATTTCCTGAAGTCCGACGGCTTCGCCAACAGAA
ACTTCATGCAGCTGATCCACGACGACAGCCTGACCTTTAAAGAGGACATCCAGA
AAGCCCAGGTGTCCGGCCAGGGCGATAGCCTGCACGAGCACATTGCCAATCT
GGCCGGCAGCCCCGCCATTAAGAAGGGCATCCTGCAGACAGTGAAGGTGGTG
GACGAGCTCGTGAAAGTGATGGGCCGGCACAAGCCCGAGAACATCGTGATCG
AAATGGCCAGAGAGAACCAGACCACCCAGAAGGGACAGAAGAACAGCCGCGA
GAGAATGAAGCGGATCGAAGAGGGCATCAAAGAGCTGGGCAGCCAGATCCTG
AAAGAACACCCCGTGGAAAACACCCAGCTGCAGAACGAGAAGCTGTACCTGTA
CTACCTGCAGAATGGGCGGGATATGTACGTGGACCAGGAACTGGACATCAACC
GGCTGTCCGACTACGATGTGGACCATATCGTGCCTCAGAGCTTTCTGAAGGAC
GACTCCATCGACAACAAGGTGCTGACCAGAAGCGACAAGAACCGGGGCAAGA
GCGACAACGTGCCCTCCGAAGAGGTCGTGAAGAAGATGAAGAACTACTGGCG
GCAGCTGCTGAACGCCAAGCTGATTACCCAGAGAAAGTTCGACAATCTGACCA
AGGCCGAGAGAGGCGGCCTGAGCGAACTGGATAAGGCCGGCTTCATCAAGAG
ACAGCTGGTGGAAACCCGGCAGATCACAAAGCACGTGGCACAGATCCTGGACT
CCCGGATGAACACTAAGTACGACGAGAATGACAAGCTGATCCGGGAAGTGAAA
GTGATCACCCTGAAGTCCAAGCTGGTGTCCGATTTCCGGAAGGATTTCCAGTTT
TACAAAGTGCGCGAGATCAACAACTACCACCACGCCCACGACGCCTACCTGAA
CGCCGTCGTGGGAACCGCCCTGATCAAAAAGTACCCTAAGCTGGAAAGCGAGT
TCGTGTACGGCGACTACAAGGTGTACGACGTGCGGAAGATGATCGCCAAGAGC
GAGCAGGAAATCGGCAAGGCTACCGCCAAGTACTTCTTCTACAGCAACATCAT
GAACTTTTTCAAGACCGAGATTACCCTGGCCAACGGCGAGATCCGGAAGCGGC
CTCTGATCGAGACAAACGGCGAAACCGGGGAGATCGTGTGGGATAAGGGCCG
GGATTTTGCCACCGTGCGGAAAGTGCTGAGCATGCCCCAAGTGAATATCGTGA
AAAAGACCGAGGTGCAGACAGGCGGCTTCAGCAAAGAGTCTATCCTGCCCAAG
AGGAACAGCGATAAGCTGATCGCCAGAAAGAAGGACTGGGACCCTAAGAAGTA
CGGCGGCTTCGACAGCCCCACCGTGGCCTATTCTGTGCTGGTGGTGGCCAAA
GTGGAAAAGGGCAAGTCCAAGAAACTGAAGAGTGTGAAAGAGCTGCTGGGGAT
CACCATCATGGAAAGAAGCAGCTTCGAGAAGAATCCCATCGACTTTCTGGAAGC
CAAGGGCTACAAAGAAGTGAAAAAGGACCTGATCATCAAGCTGCCTAAGTACTC
CCTGTTCGAGCTGGAAAACGGCCGGAAGAGAATGCTGGCCTCTGCCGGCGAA
CTGCAGAAGGGAAACGAACTGGCCCTGCCCTCCAAATATGTGAACTTCCTGTA
CCTGGCCAGCCACTATGAGAAGCTGAAGGGCTCCCCCGAGGATAATGAGCAGA
AACAGCTGTTTGTGGAACAGCACAAGCACTACCTGGACGAGATCATCGAGCAG
ATCAGCGAGTTCTCCAAGAGAGTGATCCTGGCCGACGCTAATCTGGACAAAGT
GCTGTCCGCCTACAACAAGCACCGGGATAAGCCCATCAGAGAGCAGGCCGAG
AATATCATCCACCTGTTTACCCTGACCAATCTGGGAGCCCCTGCCGCCTTCAAG
TACTTTGACACCACCATCGACCGGAAGAGGTACACCAGCACCAAAGAGGTGCT
GGACGCCACCCTGATCCACCAGAGCATCACCGGCCTGTACGAGACACGGATC
GACCTGTCTCAGCTGGGAGGTGAC_TCTGGCGGCTCAA_AAAGAACCGCCGAC
GGCAGCGAATTCGAGCCCAAGAAGAAGAGGAAAGTC_TCTGGTGGTTCTCCCA
AGAAGAAGAGGAAAGTCGGAAGCGGAGCTACTAACTTCAGCCTGCTGAAGCAG
GCTGGAGACGTGGAGGAGAACCCTGGACCTATGGTGAGCAAGGGCGAGGAGC

TGTTCACCGGGGTGGTGCCCATCCTGGTCGAGCTGGACGGCGACGTAAACGG
CCACAAGTTCAGCGTGTCCGGCGAGGGCGAGGGCGATGCCACCTACGGCAAG
CTGACCCTGAAGTTCATCTGCACCACCGGCAAGCTGCCCGTGCCCTGGCCCAC
CCTCGTGACCACCCTGACCTACGGCGTGCAGTGCTTCAGCCGCTACCCCGACC
ACATGAAGCAGCACGACTTCTTCAAGTCCGCCATGCCCGAAGGCTACGTCCAG
GAGCGCACCATCTTCTTCAAGGACGACGGCAACTACAAGACCCGCGCCGAGGT
GAAGTTCGAGGGCGACACCCTGGTGAACCGCATCGAGCTGAAGGGCATCGAC
TTCAAGGAGGACGGCAACATCCTGGGGCACAAGCTGGAGTACAACTACAACAG
CCACAACGTCTATATCATGGCCGACAAGCAGAAGAACGGCATCAAGGTGAACT
TCAAGATCCGCCACAACATCGAGGACGGCAGCGTGCAGCTCGCCGACCACTAC
CAGCAGAACACCCCCATCGGCGACGGCCCCGTGCTGCTGCCCGACAACCACT
ACCTGAGCACCCAGTCCGCCCTGAGCAAAGACCCCAACGAGAAGCGCGATCA
CATGGTCCTGCTGGAGTTCGTGACCGCCGCCGGGATCACTCTCGGCATGGAC
GAGCTGTACAAGTCTGGTGGTTCTCCCAAGAAGAAGAGGAAAGTCTAA

(b) CMV-BE4max-P2A-GFP (Addgene no. 112099)

NLS
Linker
Rat APOBEC1
Nickase SpCas9(D10A)
UGI
P2A-GFP

ATGAAACGGACAGCCGACGGAAGCGAGTTCGAGTCACCAAAGAAGAAGCGGA
AAGTC_TCCTCAGAGACTGGGCCTGTCGCCGTCGATCCAACCCTGCGCCGCCG
GATTGAACCTCACGAGTTTGAAGTGTTCTTTGACCCCCGGGAGCTGAGAAAGG
AGACATGCCTGCTGTACGAGATCAACTGGGGAGGCAGGCACTCCATCTGGAGG
CACACCTCTCAGAACACAAATAAGCACGTGGAGGTGAACTTCATCGAGAAGTTT
ACCACAGAGCGGTACTTCTGCCCCAATACCAGATGTAGCATCACATGGTTTCTG
AGCTGGTCCCCTTGCGGAGAGTGTAGCAGGGCCATCACCGAGTTCCTGTCCAG
ATATCCACACGTGACACTGTTTATCTACATCGCCAGGCTGTATCACCACGCAGA
CCCAAGGAATAGGCAGGGCCTGCGCGATCTGATCAGCTCCGGCGTGACCATC
CAGATCATGACAGAGCAGGAGTCCGGCTACTGCTGGCGGAACTTCGTGAATTA
TTCTCCTAGCAACGAGGCCCACTGGCCTAGGTACCCACACCTGTGGGTGCGCC
TGTACGTGCTGGAGCTGTATTGCATCATCCTGGGCCTGCCCCCTTGTCTGAATA
TCCTGCGGAGAAAGCAGCCCCAGCTGACCTTCTTTACAATCGCCCTGCAGTCTT
GTCACTATCAGAGGCTGCCACCCCACATCCTGTGGGCCACAGGCCTGAAG_TC
TGGAGGATCTAGCGGAGGATCCTCTGGCAGCGAGACACCAGGAACAAGCGAG
TCAGCAACACCAGAGAGCAGTGGCGGCAGCAGCGGCGGCAGC_GACAAGAAG
TACAGCATCGGCCTGGCCATCGGCACCAACTCTGTGGGCTGGGCCGTGATCAC
CGACGAGTACAAGGTGCCCAGCAAGAAATTCAAGGTGCTGGGCAACACCGACC
GGCACAGCATCAAGAAGAACCTGATCGGAGCCCTGCTGTTCGACAGCGGCGAA
ACAGCCGAGGCCACCCGGCTGAAGAGAACCGCCAGAAGAAGATACACCAGAC
GGAAGAACCGGATCTGCTATCTGCAAGAGATCTTCAGCAACGAGATGGCCAAG
GTGGACGACAGCTTCTTCCACAGACTGGAAGAGTCCTTCCTGGTGGAAGAGGA
TAAGAAGCACGAGCGGCACCCCATCTTCGGCAACATCGTGGACGAGGTGGCCT
ACCACGAGAAGTACCCCACCATCTACCACCTGAGAAAGAAACTGGTGGACAGC

ACCGACAAGGCCGACCTGCGGCTGATCTATCTGGCCCTGGCCCACATGATCAA
GTTCCGGGGCCACTTCCTGATCGAGGGCGACCTGAACCCCGACAACAGCGAC
GTGGACAAGCTGTTCATCCAGCTGGTGCAGACCTACAACCAGCTGTTCGAGGA
AAACCCCATCAACGCCAGCGGCGTGGACGCCAAGGCCATCCTGTCTGCCAGA
CTGAGCAAGAGCAGACGGCTGGAAAATCTGATCGCCCAGCTGCCCGGCGAGA
AGAAGAATGGCCTGTTCGGAAACCTGATTGCCCTGAGCCTGGGCCTGACCCCC
AACTTCAAGAGCAACTTCGACCTGGCCGAGGATGCCAAACTGCAGCTGAGCAA
GGACACCTACGACGACGACCTGGACAACCTGCTGGCCCAGATCGGCGACCAG
TACGCCGACCTGTTTCTGGCCGCCAAGAACCTGTCCGACGCCATCCTGCTGAG
CGACATCCTGAGAGTGAACACCGAGATCACCAAGGCCCCCCTGAGCGCCTCTA
TGATCAAGAGATACGACGAGCACCACCAGGACCTGACCCTGCTGAAAGCTCTC
GTGCGGCAGCAGCTGCCTGAGAAGTACAAAGAGATTTTCTTCGACCAGAGCAA
GAACGGCTACGCCGGCTACATTGACGGCGGAGCCAGCCAGGAAGAGTTCTAC
AAGTTCATCAAGCCCATCCTGGAAAAGATGGACGGCACCGAGGAACTGCTCGT
GAAGCTGAACAGAGAGGACCTGCTGCGGAAGCAGCGGACCTTCGACAACGGC
AGCATCCCCCACCAGATCCACCTGGGAGAGCTGCACGCCATTCTGCGGCGGC
AGGAAGATTTTTACCCATTCCTGAAGGACAACCGGGAAAAGATCGAGAAGATCC
TGACCTTCCGCATCCCCTACTACGTGGGCCCTCTGGCCAGGGGAAACAGCAGA
TTCGCCTGGATGACCAGAAAGAGCGAGGAAACCATCACCCCCTGGAACTTCGA
GGAAGTGGTGGACAAGGGCGCTTCCGCCCAGAGCTTCATCGAGCGGATGACC
AACTTCGATAAGAACCTGCCCAACGAGAAGGTGCTGCCCAAGCACAGCCTGCT
GTACGAGTACTTCACCGTGTATAACGAGCTGACCAAAGTGAAATACGTGACCGA
GGGAATGAGAAAGCCCGCCTTCCTGAGCGGCGAGCAGAAAAAGGCCATCGTG
GACCTGCTGTTCAAGACCAACCGGAAAGTGACCGTGAAGCAGCTGAAAGAGGA
CTACTTCAAGAAAATCGAGTGCTTCGACTCCGTGGAAATCTCCGGCGTGGAAG
ATCGGTTCAACGCCTCCCTGGGCACATACCACGATCTGCTGAAAATTATCAAGG
ACAAGGACTTCCTGGACAATGAGGAAAACGAGGACATTCTGGAAGATATCGTG
CTGACCCTGACACTGTTTGAGGACAGAGAGATGATCGAGGAACGGCTGAAAAC
CTATGCCCACCTGTTCGACGACAAAGTGATGAAGCAGCTGAAGCGGCGGAGAT
ACACCGGCTGGGGCAGGCTGAGCCGGAAGCTGATCAACGGCATCCGGGACAA
GCAGTCCGGCAAGACAATCCTGGATTTCCTGAAGTCCGACGGCTTCGCCAACA
GAAACTTCATGCAGCTGATCCACGACGACAGCCTGACCTTTAAAGAGGACATCC
AGAAAGCCCAGGTGTCCGGCCAGGGCGATAGCCTGCACGAGCACATTGCCAA
TCTGGCCGGCAGCCCCGCCATTAAGAAGGGCATCCTGCAGACAGTGAAGGTG
GTGGACGAGCTCGTGAAAGTGATGGGCCGGCACAAGCCCGAGAACATCGTGA
TCGAAATGGCCAGAGAGAACCAGACCACCCAGAAGGGACAGAAGAACAGCCG
CGAGAGAATGAAGCGGATCGAAGAGGGCATCAAAGAGCTGGGCAGCCAGATC
CTGAAAGAACACCCCGTGGAAAACACCCAGCTGCAGAACGAGAAGCTGTACCT
GTACTACCTGCAGAATGGGCGGGATATGTACGTGGACCAGGAACTGGACATCA
ACCGGCTGTCCGACTACGATGTGGACCATATCGTGCCTCAGAGCTTTCTGAAG
GACGACTCCATCGACAACAAGGTGCTGACCAGAAGCGACAAGAACCGGGGCA
AGAGCGACAACGTGCCCTCCGAAGAGGTCGTGAAGAAGATGAAGAACTACTGG
CGGCAGCTGCTGAACGCCAAGCTGATTACCCAGAGAAAGTTCGACAATCTGAC
CAAGGCCGAGAGAGGCGGCCTGAGCGAACTGGATAAGGCCGGCTTCATCAAG
AGACAGCTGGTGGAAACCCGGCAGATCACAAAGCACGTGGCACAGATCCTGGA
CTCCCGGATGAACACTAAGTACGACGAGAATGACAAGCTGATCCGGGAAGTGA
AAGTGATCACCCTGAAGTCCAAGCTGGTGTCCGATTTCCGGAAGGATTTCCAGT
TTTACAAAGTGCGCGAGATCAACAACTACCACCACGCCCACGACGCCTACCTG

AACGCCGTCGTGGGAACCGCCCTGATCAAAAAGTACCCTAAGCTGGAAAGCGA
GTTCGTGTACGGCGACTACAAGGTGTACGACGTGCGGAAGATGATCGCCAAGA
GCGAGCAGGAAATCGGCAAGGCTACCGCCAAGTACTTCTTCTACAGCAACATC
ATGAACTTTTTCAAGACCGAGATTACCCTGGCCAACGGCGAGATCCGGAAGCG
GCCTCTGATCGAGACAAACGGCGAAACCGGGGAGATCGTGTGGGATAAGGGC
CGGGATTTTGCCACCGTGCGGAAAGTGCTGAGCATGCCCCAAGTGAATATCGT
GAAAAAGACCGAGGTGCAGACAGGCGGCTTCAGCAAAGAGTCTATCCTGCCCA
AGAGGAACAGCGATAAGCTGATCGCCAGAAAGAAGGACTGGGACCCTAAGAAG
TACGGCGGCTTCGACAGCCCCACCGTGGCCTATTCTGTGCTGGTGGTGGCCAA
AGTGGAAAAGGGCAAGTCCAAGAAACTGAAGAGTGTGAAAGAGCTGCTGGGGA
TCACCATCATGGAAAGAAGCAGCTTCGAGAAGAATCCCATCGACTTTCTGGAAG
CCAAGGGCTACAAAGAAGTGAAAAAGGACCTGATCATCAAGCTGCCTAAGTACT
CCCTGTTCGAGCTGGAAAACGGCCGGAAGAGAATGCTGGCCTCTGCCGGCGA
ACTGCAGAAGGGAAACGAACTGGCCCTGCCCTCCAAATATGTGAACTTCCTGTA
CCTGGCCAGCCACTATGAGAAGCTGAAGGGCTCCCCCGAGGATAATGAGCAGA
AACAGCTGTTTGTGGAACAGCACAAGCACTACCTGGACGAGATCATCGAGCAG
ATCAGCGAGTTCTCCAAGAGAGTGATCCTGGCCGACGCTAATCTGGACAAAGT
GCTGTCCGCCTACAACAAGCACCGGGATAAGCCCATCAGAGAGCAGGCCGAG
AATATCATCCACCTGTTTACCCTGACCAATCTGGGAGCCCCTGCCGCCTTCAAG
TACTTTGACACCACCATCGACCGGAAGAGGTACACCAGCACCAAAGAGGTGCT
GGACGCCACCCTGATCCACCAGAGCATCACCGGCCTGTACGAGACACGGATC
GACCTGTCTCAGCTGGGAGGTGAC_AGCGGCGGGAGCGGCGGGAGCGGGGG
GAGC_ACTAATCTGAGCGACATCATTGAGAAGGAGACTGGGAAACAGCTGGTC
ATTCAGGAGTCCATCCTGATGCTGCCTGAGGAGGTGGAGGAAGTGATCGGCAA
CAAGCCAGAGTCTGACATCCTGGTGCACACCGCCTACGACGAGTCCACAGATG
AGAATGTGATGCTGCTGACCTCTGACGCCCCCGAGTATAAGCCTTGGGCCCTG
GTCATCCAGGATTCTAACGGCGAGAATAAGATCAAGATGCTG_AGCGGAGGAT
CCGGAGGATCTGGAGGCAGC_ACCAACCTGTCTGACATCATCGAGAAGGAGAC
AGGCAAGCAGCTGGTCATCCAGGAGAGCATCCTGATGCTGCCCGAAGAAGTCG
AAGAAGTGATCGGAAACAAGCCTGAGAGCGATATCCTGGTCCATACCGCCTAC
GACGAGAGTACCGACGAAAATGTGATGCTGCTGACATCCGACGCCCCAGAGTA
TAAGCCCTGGGCTCTGGTCATCCAGGATTCCAACGGAGAGAACAAAATCAAAAT
GCTG_TCTGGCGGCTCA_AAAAGAACCGCCGACGGCAGCGAATTCGAGCCCAA
GAAGAAGAGGAAAGTC_TCTGGTGGTTCTCCCAAGAAGAAGAGGAAAGTCGGA
AGCGGAGCTACTAACTTCAGCCTGCTGAAGCAGGCTGGAGACGTGGAGGAGA
ACCCTGGACCTATGGTGAGCAAGGGCGAGGAGCTGTTCACCGGGGTGGTGCC
CATCCTGGTCGAGCTGGACGGCGACGTAAACGGCCACAAGTTCAGCGTGTCC
GGCGAGGGCGAGGGCGATGCCACCTACGGCAAGCTGACCCTGAAGTTCATCT
GCACCACCGGCAAGCTGCCCGTGCCCTGGCCCACCCTCGTGACCACCCTGAC
CTACGGCGTGCAGTGCTTCAGCCGCTACCCCGACCACATGAAGCAGCACGACT
TCTTCAAGTCCGCCATGCCCGAAGGCTACGTCCAGGAGCGCACCATCTTCTTC
AAGGACGACGGCAACTACAAGACCCGCGCCGAGGTGAAGTTCGAGGGCGACA
CCCTGGTGAACCGCATCGAGCTGAAGGGCATCGACTTCAAGGAGGACGGCAA
CATCCTGGGGCACAAGCTGGAGTACAACTACAACAGCCACAACGTCTATATCAT
GGCCGACAAGCAGAAGAACGGCATCAAGGTGAACTTCAAGATCCGCCACAACA
TCGAGGACGGCAGCGTGCAGCTCGCCGACCACTACCAGCAGAACACCCCCAT
CGGCGACGGCCCCGTGCTGCTGCCCGACAACCACTACCTGAGCACCCAGTCC
GCCCTGAGCAAAGACCCCAACGAGAAGCGCGATCACATGGTCCTGCTGGAGTT

CGTGACCGCCGCCGGGATCACTCTCGGCATGGACGAGCTGTACAAGTCTGGT
GGTTCTCCCAAGAAGAAGAGGAAAGTCTAA

(c) CMV-ABE8e (Addgene no. 138489)

NLS
Linker
ecTadA(8e)
Nickase SpCas9(D10A)

ATGAAACGGACAGCCGACGGAAGCGAGTTCGAGTCACCAAAGAAGAAGCGGA
AAGTC_TCTGAGGTGGAGTTTTCCCACGAGTACTGGATGAGACATGCCCTGACC
CTGGCCAAGAGGGCACGGGATGAGAGGGAGGTGCCTGTGGGAGCCGTGCTG
GTGCTGAACAATAGAGTGATCGGCGAGGGCTGGAACAGAGCCATCGGCCTGC
ACGACCCAACAGCCCATGCCGAAATTATGGCCCTGAGACAGGGCGGCCTGGT
CATGCAGAACTACAGACTGATTGACGCCACCCTGTACGTGACATTCGAGCCTTG
CGTGATGTGCGCCGGCGCCATGATCCACTCTAGGATCGGCCGCGTGGTGTTTG
GCGTGAGGAACTCAAAAAGAGGCGCCGCAGGCTCCCTGATGAACGTGCTGAA
CTACCCCGGCATGAATCACCGCGTCGAAATTACCGAGGGAATCCTGGCAGATG
AATGTGCCGCCCTGCTGTGCGATTTCTATCGGATGCCTAGACAGGTGTTCAATG
CTCAGAAGAAGGCCCAGAGCTCCATCAAC_TCCGGAGGATCTAGCGGAGGCTC
CTCTGGCTCTGAGACACCTGGCACAAGCGAGAGCGCAACACCTGAAAGCAGC
GGGGGCAGCAGCGGGGGGTCA_GACAAGAAGTACAGCATCGGCCTGGCCATC
GGCACCAACTCTGTGGGCTGGGCCGTGATCACCGACGAGTACAAGGTGCCCA
GCAAGAAATTCAAGGTGCTGGGCAACACCGACCGGCACAGCATCAAGAAGAAC
CTGATCGGAGCCCTGCTGTTCGACAGCGGCGAAACAGCCGAGGCCACCCGGC
TGAAGAGAACCGCCAGAAGAAGATACACCAGACGGAAGAACCGGATCTGCTAT
CTGCAAGAGATCTTCAGCAACGAGATGGCCAAGGTGGACGACAGCTTCTTCCA
CAGACTGGAAGAGTCCTTCCTGGTGGAAGAGGATAAGAAGCACGAGCGGCAC
CCCATCTTCGGCAACATCGTGGACGAGGTGGCCTACCACGAGAAGTACCCCAC
CATCTACCACCTGAGAAAGAAACTGGTGGACAGCACCGACAAGGCCGACCTGC
GGCTGATCTATCTGGCCCTGGCCCACATGATCAAGTTCCGGGGCCACTTCCTG
ATCGAGGGCGACCTGAACCCCGACAACAGCGACGTGGACAAGCTGTTCATCCA
GCTGGTGCAGACCTACAACCAGCTGTTCGAGGAAAACCCCATCAACGCCAGCG
GCGTGGACGCCAAGGCCATCCTGTCTGCCAGACTGAGCAAGAGCAGACGGCT
GGAAAATCTGATCGCCCAGCTGCCCGGCGAGAAGAAGAATGGCCTGTTCGGAA
ACCTGATTGCCCTGAGCCTGGGCCTGACCCCCAACTTCAAGAGCAACTTCGAC
CTGGCCGAGGATGCCAAACTGCAGCTGAGCAAGGACACCTACGACGACGACC
TGGACAACCTGCTGGCCCAGATCGGCGACCAGTACGCCGACCTGTTTCTGGCC
GCCAAGAACCTGTCCGACGCCATCCTGCTGAGCGACATCCTGAGAGTGAACAC
CGAGATCACCAAGGCCCCCCTGAGCGCCTCTATGATCAAGAGATACGACGAGC
ACCACCAGGACCTGACCCTGCTGAAAGCTCTCGTGCGGCAGCAGCTGCCTGA
GAAGTACAAAGAGATTTTCTTCGACCAGAGCAAGAACGGCTACGCCGGCTACA
TTGACGGCGGAGCCAGCCAGGAAGAGTTCTACAAGTTCATCAAGCCCATCCTG
GAAAAGATGGACGGCACCGAGGAACTGCTCGTGAAGCTGAACAGAGAGGACC
TGCTGCGGAAGCAGCGGACCTTCGACAACGGCAGCATCCCCCACCAGATCCA
CCTGGGAGAGCTGCACGCCATTCTGCGGCGGCAGGAAGATTTTTACCCATTCC

TGAAGGACAACCGGGAAAAGATCGAGAAGATCCTGACCTTCCGCATCCCCTAC
TACGTGGGCCCTCTGGCCAGGGGAAACAGCAGATTCGCCTGGATGACCAGAAA
GAGCGAGGAAACCATCACCCCCTGGAACTTCGAGGAAGTGGTGGACAAGGGC
GCTTCCGCCCAGAGCTTCATCGAGCGGATGACCAACTTCGATAAGAACCTGCC
CAACGAGAAGGTGCTGCCCAAGCACAGCCTGCTGTACGAGTACTTCACCGTGT
ATAACGAGCTGACCAAAGTGAAATACGTGACCGAGGGAATGAGAAAGCCCGCC
TTCCTGAGCGGCGAGCAGAAAAAGGCCATCGTGGACCTGCTGTTCAAGACCAA
CCGGAAAGTGACCGTGAAGCAGCTGAAAGAGGACTACTTCAAGAAAATCGAGT
GCTTCGACTCCGTGGAAATCTCCGGCGTGGAAGATCGGTTCAACGCCTCCCTG
GGCACATACCACGATCTGCTGAAAATTATCAAGGACAAGGACTTCCTGGACAAT
GAGGAAAACGAGGACATTCTGGAAGATATCGTGCTGACCCTGACACTGTTTGA
GGACAGAGAGATGATCGAGGAACGGCTGAAAACCTATGCCCACCTGTTCGACG
ACAAAGTGATGAAGCAGCTGAAGCGGCGGAGATACACCGGCTGGGGCAGGCT
GAGCCGGAAGCTGATCAACGGCATCCGGGACAAGCAGTCCGGCAAGACAATC
CTGGATTTCCTGAAGTCCGACGGCTTCGCCAACAGAAACTTCATGCAGCTGATC
CACGACGACAGCCTGACCTTTAAAGAGGACATCCAGAAAGCCCAGGTGTCCGG
CCAGGGCGATAGCCTGCACGAGCACATTGCCAATCTGGCCGGCAGCCCCGCC
ATTAAGAAGGGCATCCTGCAGACAGTGAAGGTGGTGGACGAGCTCGTGAAAGT
GATGGGCCGGCACAAGCCCGAGAACATCGTGATCGAAATGGCCAGAGAGAAC
CAGACCACCCAGAAGGGACAGAAGAACAGCCGCGAGAGAATGAAGCGGATCG
AAGAGGGCATCAAAGAGCTGGGCAGCCAGATCCTGAAAGAACACCCCGTGGAA
AACACCCAGCTGCAGAACGAGAAGCTGTACCTGTACTACCTGCAGAATGGGCG
GGATATGTACGTGGACCAGGAACTGGACATCAACCGGCTGTCCGACTACGATG
TGGACCATATCGTGCCTCAGAGCTTTCTGAAGGACGACTCCATCGACAACAAG
GTGCTGACCAGAAGCGACAAGAACCGGGGCAAGAGCGACAACGTGCCCTCCG
AAGAGGTCGTGAAGAAGATGAAGAACTACTGGCGGCAGCTGCTGAACGCCAAG
CTGATTACCCAGAGAAAGTTCGACAATCTGACCAAGGCCGAGAGAGGCGGCCT
GAGCGAACTGGATAAGGCCGGCTTCATCAAGAGACAGCTGGTGGAAACCCGG
CAGATCACAAAGCACGTGGCACAGATCCTGGACTCCCGGATGAACACTAAGTA
CGACGAGAATGACAAGCTGATCCGGGAAGTGAAAGTGATCACCCTGAAGTCCA
AGCTGGTGTCCGATTTCCGGAAGGATTTCCAGTTTTACAAAGTGCGCGAGATCA
ACAACTACCACCACGCCCACGACGCCTACCTGAACGCCGTCGTGGGAACCGC
CCTGATCAAAAAGTACCCTAAGCTGGAAAGCGAGTTCGTGTACGGCGACTACA
AGGTGTACGACGTGCGGAAGATGATCGCCAAGAGCGAGCAGGAAATCGGCAA
GGCTACCGCCAAGTACTTCTTCTACAGCAACATCATGAACTTTTTCAAGACCGA
GATTACCCTGGCCAACGGCGAGATCCGGAAGCGGCCTCTGATCGAGACAAAC
GGCGAAACCGGGGAGATCGTGTGGGATAAGGGCCGGGATTTTGCCACCGTGC
GGAAAGTGCTGAGCATGCCCCAAGTGAATATCGTGAAAAAGACCGAGGTGCAG
ACAGGCGGCTTCAGCAAAGAGTCTATCCTGCCCAAGAGGAACAGCGATAAGCT
GATCGCCAGAAAGAAGGACTGGGACCCTAAGAAGTACGGCGGCTTCGACAGC
CCCACCGTGGCCTATTCTGTGCTGGTGGTGGCCAAAGTGGAAAAGGGCAAGTC
CAAGAAACTGAAGAGTGTGAAAGAGCTGCTGGGGATCACCATCATGGAAAGAA
GCAGCTTCGAGAAGAATCCCATCGACTTTCTGGAAGCCAAGGGCTACAAAGAA
GTGAAAAAGGACCTGATCATCAAGCTGCCTAAGTACTCCCTGTTCGAGCTGGAA
AACGGCCGGAAGAGAATGCTGGCCTCTGCCGGCGAACTGCAGAAGGGAAACG
AACTGGCCCTGCCCTCCAAATATGTGAACTTCCTGTACCTGGCCAGCCACTATG
AGAAGCTGAAGGGCTCCCCCGAGGATAATGAGCAGAAACAGCTGTTTGTGGAA
CAGCACAAGCACTACCTGGACGAGATCATCGAGCAGATCAGCGAGTTCTCCAA

GAGAGTGATCCTGGCCGACGCTAATCTGGACAAAGTGCTGTCCGCCTACAACA
AGCACCGGGATAAGCCCATCAGAGAGCAGGCCGAGAATATCATCCACCTGTTT
ACCCTGACCAATCTGGGAGCCCCTGCCGCCTTCAAGTACTTTGACACCACCAT
CGACCGGAAGAGGTACACCAGCACCAAAGAGGTGCTGGACGCCACCCTGATC
CACCAGAGCATCACCGGCCTGTACGAGACACGGATCGACCTGTCTCAGCTGGG
AGGTGAC_TCTGGCGGCTCAA_AAAGAACCGCCGACGGCAGCGAATTCGAGCC
CAAGAAGAAGAGGAAAGTCTAA

(d) CAG-nSpCas9-pmCDA1-P2A-GFP (Target-AID; Addgene no. 131300)

NLS
Linker
Nickase SpCas9(D10A)
3xFLAG
pmCDA1
UGI
P2A-EGFP

ATGGCACCGAAGAAGAAGCGTAAAGTCGGAATCCACGGAGTTCCTGCGGCA_A
TGGACAAGAAGTACTCCATTGGGCTCGCTATCGGCACAAACAGCGTCGGTTGG
GCCGTCATTACGGACGAGTACAAGGTGCCGAGCAAAAAATTCAAAGTTCTGGG
CAATACCGATCGCCACAGCATAAAGAAGAACCTCATTGGCGCCCTCCTGTTCGA
CTCCGGGGAGACGGCCGAAGCCACGCGGCTCAAAAGAACAGCACGGCGCAGA
TATACCCGCAGAAAGAATCGGATCTGCTACCTGCAGGAGATCTTTAGTAATGAG
ATGGCTAAGGTGGATGACTCTTTCTTCCATAGGCTGGAGGAGTCCTTTTTGGTG
GAGGAGGATAAAAAGCACGAGCGCCACCCAATCTTTGGCAATATCGTGGACGA
GGTGGCGTACCATGAAAAGTACCCAACCATATATCATCTGAGGAAGAAGCTTGT
AGACAGTACTGATAAGGCTGACTTGCGGTTGATCTATCTCGCGCTGGCGCATAT
GATCAAATTTCGGGGACACTTCCTCATCGAGGGGGACCTGAACCCAGACAACA
GCGATGTCGACAAACTCTTTATCCAACTGGTTCAGACTTACAATCAGCTTTTCGA
AGAGAACCCGATCAACGCATCCGGAGTTGACGCCAAAGCAATCCTGAGCGCTA
GGCTGTCCAAATCCCGGCGGCTCGAAAACCTCATCGCACAGCTCCCTGGGGA
GAAGAAGAACGGCCTGTTTGGTAATCTTATCGCCCTGTCACTCGGGCTGACCC
CCAACTTTAAATCTAACTTCGACCTGGCCGAAGATGCCAAGCTTCAACTGAGCA
AAGACACCTACGATGATGATCTCGACAATCTGCTGGCCCAGATCGGCGACCAG
TACGCAGACCTTTTTTTGGCGGCAAAGAACCTGTCAGACGCCATTCTGCTGAGT
GATATTCTGCGAGTGAACACGGAGATCACCAAAGCTCCGCTGAGCGCTAGTAT
GATCAAGCGCTATGATGAGCACCACCAAGACTTGACTTTGCTGAAGGCCCTTGT
CAGACAGCAACTGCCTGAGAAGTACAAGGAAATTTTCTTCGATCAGTCTAAAAA
TGGCTACGCCGGATACATTGACGGCGGAGCAAGCCAGGAGGAATTTTACAAAT
TTATTAAGCCCATCTTGGAAAAAATGGACGGCACCGAGGAGCTGCTGGTAAAG
CTTAACAGAGAAGATCTGTTGCGCAAACAGCGCACTTTCGACAATGGAAGCATC
CCCCACCAGATTCACCTGGGCGAACTGCACGCTATCCTCAGGCGGCAAGAGGA
TTTCTACCCCTTTTTGAAAGATAACAGGGAAAAGATTGAGAAAATCCTCACATTT
CGGATACCCTACTATGTAGGCCCCCTCGCCCGGGGAAATTCCAGATTCGCGTG
GATGACTCGCAAATCAGAAGAGACCATCACTCCCTGGAACTTCGAGGAAGTCG
TGGATAAGGGGGCCTCTGCCCAGTCCTTCATCGAAAGGATGACTAACTTTGATA
AAAATCTGCCTAACGAAAAGGTGCTTCCTAAACACTCTCTGCTGTACGAGTACT

TCACAGTTTATAACGAGCTCACCAAGGTCAAATACGTCACAGAAGGGATGAGAA
AGCCAGCATTCCTGTCTGGAGAGCAGAAGAAAGCTATCGTGGACCTCCTCTTC
AAGACGAACCGGAAAGTTACCGTGAAACAGCTCAAAGAAGACTATTTCAAAAAG
ATTGAATGTTTCGACTCTGTTGAAATCAGCGGAGTGGAGGATCGCTTCAACGCA
TCCCTGGGAACGTATCACGATCTCCTGAAAATCATTAAAGACAAGGACTTCCTG
GACAATGAGGAGAACGAGGACATTCTTGAGGACATTGTCCTCACCCTTACGTTG
TTTGAAGATAGGGAGATGATTGAAGAACGCTTGAAAACTTACGCTCATCTCTTC
GACGACAAAGTCATGAAACAGCTCAAGAGGCGCCGATATACAGGATGGGGGC
GGCTGTCAAGAAAACTGATCAATGGGATCCGAGACAAGCAGAGTGGAAAGACA
ATCCTGGATTTTCTTAAGTCCGATGGATTTGCCAACCGGAACTTCATGCAGTTG
ATCCATGATGACTCTCTCACCTTTAAGGAGGACATCCAGAAAGCACAAGTTTCT
GGCCAGGGGGACAGTCTTCACGAGCACATCGCTAATCTTGCAGGTAGCCCAGC
TATCAAAAAGGGAATACTGCAGACCGTTAAGGTCGTGGATGAACTCGTCAAAGT
AATGGGAAGGCATAAGCCCGAGAATATCGTTATCGAGATGGCCCGAGAGAACC
AAACTACCCAGAAGGGACAGAAGAACAGTAGGGAAAGGATGAAGAGGATTGAA
GAGGGTATAAAAGAACTGGGGTCCCAAATCCTTAAGGAACACCCAGTTGAAAAC
ACCCAGCTTCAGAATGAGAAGCTCTACCTGTACTACCTGCAGAACGGCAGGGA
CATGTACGTGGATCAGGAACTGGACATCAATCGGCTCTCCGACTACGACGTGG
ATCATATCGTGCCCCAGTCTTTTCTCAAAGATGATTCTATTGATAATAAAGTGTT
GACAAGATCCGATAAAAATAGAGGGAAGAGTGATAACGTCCCCTCAGAAGAAG
TTGTCAAGAAAATGAAAAATTATTGGCGGCAGCTGCTGAACGCCAAACTGATCA
CACAACGGAAGTTCGATAATCTGACTAAGGCTGAACGAGGTGGCCTGTCTGAG
TTGGATAAAGCCGGCTTCATCAAAAGGCAGCTTGTTGAGACACGCCAGATCAC
CAAGCACGTGGCCCAAATTCTCGATTCACGCATGAACACCAAGTACGATGAAAA
TGACAAACTGATTCGAGAGGTGAAAGTTATTACTCTGAAGTCTAAGCTGGTCTC
AGATTTCAGAAAGGACTTTCAGTTTTATAAGGTGAGAGAGATCAACAATTACCAC
CATGCGCATGATGCCTACCTGAATGCAGTGGTAGGCACTGCACTTATCAAAAAA
TATCCCAAGCTTGAATCTGAATTTGTTTACGGAGACTATAAAGTGTACGATGTTA
GGAAAATGATCGCAAAGTCTGAGCAGGAAATAGGCAAGGCCACCGCTAAGTAC
TTCTTTTACAGCAATATTATGAATTTTTTCAAGACCGAGATTACACTGGCCAATG
GAGAGATTCGGAAGCGACCACTTATCGAAACAAACGGAGAAACAGGAGAAATC
GTGTGGGACAAGGGTAGGGATTTCGCGACAGTCCGGAAGGTCCTGTCCATGC
CGCAGGTGAACATCGTTAAAAAGACCGAAGTACAGACCGGAGGCTTCTCCAAG
GAAAGTATCCTCCCGAAAAGGAACAGCGACAAGCTGATCGCACGCAAAAAAGA
TTGGGACCCCAAGAAATACGGCGGATTCGATTCTCCTACAGTCGCTTACAGTGT
ACTGGTTGTGGCCAAAGTGGAGAAAGGGAAGTCTAAAAAACTCAAAAGCGTCA
AGGAACTGCTGGGCATCACAATCATGGAGCGATCAAGCTTCGAAAAAAACCCC
ATCGACTTTCTCGAGGCGAAAGGATATAAAGAGGTCAAAAAAGACCTCATCATT
AAGCTTCCCAAGTACTCTCTCTTTGAGCTTGAAAACGGCCGGAAACGAATGCTC
GCTAGTGCGGGCGAGCTGCAGAAAGGTAACGAGCTGGCACTGCCCTCTAAATA
CGTTAATTTCTTGTATCTGGCCAGCCACTATGAAAAGCTCAAAGGGTCTCCCGA
AGATAATGAGCAGAAGCAGCTGTTCGTGGAACAACACAAACACTACCTTGATGA
GATCATCGAGCAAATAAGCGAATTCTCCAAAAGAGTGATCCTCGCCGACGCTAA
CCTCGATAAGGTGCTTTCTGCTTACAATAAGCACAGGGATAAGCCCATCAGGGA
GCAGGCAGAAAACATTATCCACTTGTTTACTCTGACCAACTTGGGCGCGCCTGC
AGCCTTCAAGTACTTCGACACCACCATAGACAGAAAGCGGTACACCTCTACAAA
GGAGGTCCTGGACGCCACACTGATTCATCAGTCAATTACGGGGCTCTATGAAA
CAAGAATCGACCTCTCTCAGCTCGGTGGAGAC_AGCAGGGCTGAC_CCCAAGA

AGAAGAGGAAGGTG_GGTGGAGGAGGTACCGGCGGTGGAGGCTCAGCAGAAT
ACGTACGAGCTCTGTTTGACTTCAATGGGAATGACGAGGAGGATCTCCCCTTTA
AGAAGGGCGATATTCTCCGCATCAGAGATAAGCCCGAAGAACAATGGTGGAAT
GCCGAGGATAGCGAAGGGAAAAGGGGCATGATTCTGGTGCCATATGTGGAGAA
ATATTCCGGT_GACTACAAAGACCATGATGGGGATTACAAAGACCACGACATCG
ACTACAAAGACGACGACGATAAA_TCAGGGATGACAGACGCCGAGTACGTGCG
CATTCATGAGAAACTGGATATTTACACCTTCAAGAAGCAGTTCTTCAACAACAAG
AAATCTGTGTCACACCGCTGCTACGTGCTGTTTGAGTTGAAGCGAAGGGGCGA
AAGAAGGGCTTGCTTTTGGGGCTATGCCGTCAACAAGCCCCAAAGTGGCACCG
AGAGAGGAATACACGCTGAGATATTCAGTATCCGAAAGGTGGAAGAGTATCTTC
GGGATAATCCT_GGGCAGTTTACGATCAACTGGTATTCCAGCTGGAGTCCTTGC
GCTGATTGTGCCGAGAAAATTCTGGAATGGTATAATCAGGAACTTCGGGGAAAC
GGGCACACATTGAAAATCTGGGCCTGCAAGCTGTACTACGAGAAGAATGCCCG
GAACCAGATAGGACTCTGGAATCTGAGGGACAATGGTGTAGGCCTGAACGTGA
TGGTTTCCGAGCACTATCAGTGTTGTCGGAAGATTTTCATCCAAAGCTCTCATAA
CCAGCTCAATGAAAACCGCTGGTTGGAGAAAACACTGAAACGTGCGGAGAAGT
GGAGATCCGAGCTGAGCATCATGATCCAGGTCAAGATTCTGCATACCACTAAGT
CTCCAGCCGTTGGT_CCCAAGAAGAAAAGAAAAGTC_GGTACC_ATGACCAACC
TTTCCGACATCATAGAGAAGGAAACAGGCAAACAGTTGGTCATCCAAGAGTCGA
TACTCATGCTTCCTGAAGAAGTTGAGGAGGTCATTGGGAATAAGCCGGAAAGT
GACATTCTCGTACACACTGCGTATGATGAGAGCACCGATGAGAACGTGATGCT
GCTCACGTCAGATGCCCCAGAGTACAAACCCTGGGCTCTGGTGATTCAGGACT
CTAATGGAGAGAACAAGATCAAGATGCTA_GGAGGCGGTGGAAGCGGCGCAAC
AAACTTCTCTCTGCTGAAACAAGCCGGAGATGTCGAAGAGAATCCTGGACCGAT
GGTGAGCAAGGGCGAGGAGCTGTTCACCGGGGTGGTGCCCATCCTGGTCGAG
CTGGACGGCGACGTAAACGGCCACAAGTTCAGCGTGTCCGGCGAGGGCGAGG
GCGATGCCACCTACGGCAAGCTGACCCTGAAGTTCATCTGCACCACCGGCAAG
CTGCCCGTGCCCTGGCCCACCCTCGTGACCACCCTGACCTACGGCGTGCAGT
GCTTCAGCCGCTACCCCGACCACATGAAGCAGCACGACTTCTTCAAGTCCGCC
ATGCCCGAAGGCTACGTCCAGGAGCGCACCATCTTCTTCAAGGACGACGGCAA
CTACAAGACCCGCGCCGAGGTGAAGTTCGAGGGCGACACCCTGGTGAACCGC
ATCGAGCTGAAGGGCATCGACTTCAAGGAGGACGGCAACATCCTGGGGCACAA
GCTGGAGTACAACTACAACAGCCACAACGTCTATATCATGGCCGACAAGCAGA
AGAACGGCATCAAGGTGAACTTCAAGATCCGCCACAACATCGAGGACGGCAGC
GTGCAGCTCGCCGACCACTACCAGCAGAACACCCCCATCGGCGACGGCCCCG
TGCTGCTGCCCGACAACCACTACCTGAGCACCCAGTCCGCCCTGAGCAAAGAC
CCCAACGAGAAGCGCGATCACATGGTCCTGCTGGAGTTCGTGACCGCCGCCG
GGATCACTCTCGGCATGGACGAGCTGTACAAGTGA
